# *ZNF827* pleiotropic cardiovascular risk locus involves regulation by Nuclear factor-1

**DOI:** 10.1101/2024.01.23.576845

**Authors:** Yingwei Liu, Lu Liu, Asraa Esmael, Margaux-Alison Fustier, Adrien Georges, Nabila Bouatia-Naji

## Abstract

**Introduction:** SCAD (spontaneous coronary artery dissection) is a form of myocardial infarction that disproportionately affects young women. SCAD is caused by the formation of a hematoma within coronary artery wall, resulting in its blockage. Sixteen SCAD susceptibility loci are currently known, including *ZNF827* gene locus on chromosome 4. Several common genetic variations within *ZNF827* are also linked to coronary artery disease (CAD), systolic blood pressure (SBP), and ascending aortic diameter. The molecular processes driving these genetic connections, however, are unknown.

**Methods:** We performed fine-mapping and genetic colocalization analysis at *ZNF827* locus on multiple GWAS signals and quantitative expression traits. We explored the regulatory function of sequences of interest using a luciferase reporter system in rat smooth muscle cells. We used siRNA to knockdown the expression of gene and assessed transcriptomic changes through bulk RNA-Seq.

**Results:** The genetic association signals for SCAD, CAD, blood pressure, and aorta dimensions could all be explained by a single causal variant. We identified rs13128814 as the most likely causative variant. This variant overlaps epigenetic regulatory markers specifically active in vascular SMCs and fibroblasts. A 1kb region containing the SCAD-risk allele (A) of rs13128814 variant had a transcriptional activation impact, while the protective allele (G) had no effect. *In silico* predictions found nuclear factor 1 (NF1) transcription factors preferentially bind to the A allele of rs13128814. We found that overexpressing NFIA (Nuclear Factor 1 A-Type) activated transcription in this area. SCAD association and *ZNF827* eQTL colocalization in various vascular tissues suggested the latter as a plausible target gene in this locus. Knockdown of ZNF827 in iPSC-derived SMCs and fibroblasts, followed by RNA-seq, identified a large number of dysregulated genes, including multiple genes involved in cardiovascular disease.

**Conclusion:** Our findings in *ZNF827* locus support the hypothesis that the rs13128814 variant, which may affect the binding of NF1 family transcription factors and cause *ZNF827* expression to be deregulated, is the molecular mechanism underpinning the relationship of the *ZNF827* locus with numerous cardiovascular disorders. *ZNF827* acts as a regulator of gene expression in vascular SMCs and fibroblasts. Further investigation of *ZNF827* function in vascular tissue will be required to understand the implications of its dysregulation on cardiovascular physiopathology.

## Introduction

Spontaneous coronary artery dissection (SCAD) is an acute coronary syndrome primarily affecting young women, representing up to 35% of acute coronary syndromes in women under the age of 50^1,2^. Unlike atherosclerosis-related coronary artery disease (CAD), SCAD results from a hematoma obstructing the coronary artery, causing arterial dissection^1^. Despite substantial clinical research, the mechanisms of SCAD remain obscure. While traditional risk factors like age, smoking, high blood pressure, and high cholesterol levels are associated with myocardial infarction (MI) and its most frequent cause, coronary artery disease (CAD) ^3,4^, SCAD primarily affects younger, often healthy individuals, challenging the applicability of these conventional risk factors ^5,6^. This has motivated the investigation of alternative causes, such as genetic predisposition, to understand the mechanisms at stake in SCAD.

We recently performed the largest meta-analysis of SCAD genome-wide association studies (GWAS), including a total of 1917 SCAD patients and 9232 controls from 8 independent case-control studies^7^. We identified a total of 16 genetic risk loci, of which 11 were newly discovered, and found potential links to multiple genes involved in extracellular matrix synthesis and maintenance, SMC contraction and tissue-mediated coagulation. However, all identified variants were non-coding, and causal variants and target genes remain to be unambiguously identified at most SCAD *loci*. A particularly striking observation is the extent of the genetic basis of SCAD that is shared with other cardiovascular traits and diseases. For example, three of SCAD loci are consistently associated to fibromuscular dysplasia (FMD), a non-atherosclerotic systemic arteriopathy frequently found in SCAD patients ^2,8^, and migraine, a common co-morbidity of SCAD patients^9^. On the other hand, about half of SCAD loci are at least suggestively associated to CAD, and in every case the risk allele of SCAD is a protective allele for CAD^7^. This relation was verified at the genomic level where an inverse genetic correlation was identified between SCAD and CAD^7^.

Among newly identified SCAD genetic risk loci, a locus close to *Zing Finger Protein 827* (*ZNF827*) appears as quite paradigmatic. Indeed, this locus was identified as associated, at least suggestively, to multiple cardiovascular traits and diseases^7^, with an apparent inverse association between SCAD and CAD. On the other hand, the regulatory mechanisms and potentially relevant target gene(s) at this locus remain elusive. SCAD lead SNP is located within an intronic region of *ZNF827*, encodes a protein predicted to bind nucleic acids, but its potential functions in the context cardiovascular pathologies are unknown. Interestingly, several studies pointed to potential functions of *ZNF827* acting as a major regulator both through its RNA and protein products ^10–12^.

In the present study, we aim at clarifying the potential mechanisms linking genetic variants at *ZNF827* locus with several cardiovascular diseases. First, we aimed at identifying all cardiovascular traits and diseases which could share a common risk variant at *ZNF827* locus, and we used multi-trait colocalization to identify this potential causal variant. Second, we tried to determine how this variant could alter transcriptional regulation. Third, we aimed to unambiguously determine the target gene(s) at *ZNF827* locus. Finally, we examined the transcriptional effect of *ZNF827* knockdown in vascular smooth muscle cells and fibroblasts. Altogether, our results provide an exploration of regulatory mechanisms at a highly pleiotropic genetic locus involved in vascular diseases.

## Methods

### Genetic lookup and colocalization

Proxies of SCAD lead SNP rs1507928 were retrieved from 1000 genome reference panel using LDProxy function of LDLinkR package (v1.3.0), using European populations and GRCh37 genome build. SNPs with r^2^ > 0.7 were used to probe GWAS Catalog database (downloaded on 2023 October 11^th^). Traits with genome-wide association (*P_leadSNP_* < 5×10^-8^) and direct cardiovascular relevance were retained for further analyses. Summary statistics were retrieved from individual studies ^7,13–17^. Multitrait colocalization was performed using HyPrColoc package (v1.0)^18^ with default options. All traits clustered together, and the global posterior probability for these traits to share a single causal variant was reported. SNP score (posterior probability of being causal) was also used to identify potential causal SNPs. For colocalization of SCAD association with *ZNF827* eQTL, signal colocalization was evaluated using R coloc package (v5.1.0) with default values as priors^19^. eQTL associations for all variants were retrieved from GTEx website (v8, gtexportal.org). The H4 coefficient indicating the probability of the two traits to share a causal variant was reported. Colocalization plot was generated using locuscomparer package (v1.0.0).

### Epigenomic data

Single nucleus ATAC-Seq in coronary arteries were retrieved from Sequence Read Archive (GSE175621)^20^. Clustering and cell type attribution were performed following the pipeline provided by authors using ArchR package (v1.0.1). Fragments attributed to each cell type were retrieved in bed format, converted to bam format using bedtools BedtoBam (v2.3.0) and bigwig coverage files were generated using deeptools bamcoverage (v3.5.4). Coronary Artery RNA-Seq dataset (raw counts) was retrieved from GTEx website. Read normalization was performed using DESeq2 package (v1.38.3). Coronary Artery Histone ChIP was retrieved from ENCODE (H3K4me3: ENCFF811RQX, H3K27Ac: ENCFF130NUG). Coronary Artery ATAC -Seq, Human Dermal Fibroblast and Coronary Artery SMC were retrieved from Sequence Read Archive (Bioprojects PRJNA295524 and PRJNA69000) and analyzed as previously described^13^. RNA-Seq and ATAC-Seq datasets during iPSC differentiation into SMC were retrieved from Sequence Read Archive (Bioproject PRJNA899672) and analyzed as previously described^21^.

### Cell culture

Rat smooth muscle cells (A7r5) were acquired from ATCC® in Manassas, Virginia, and grown in DMEM with the addition of 10% FBS from Thermo Fisher Scientific in Waltham, Massachusetts, USA. BJ fibroblasts were acquired from ATCC® in Manassas, Virginia, and grown in DMEM with the addition of 10% FBS from Thermo Fisher Scientific in Waltham, Massachusetts, USA. Human iPSC line SKiPS-31.3 was obtained by reprogramming of human dermal fibroblast of a healthy male adult volunteer as previously described^22^. iPSC lines 11.10 and 12.10 were purchased from Cell Applications (San Diego, CA). All iPSC lines were nurtured in mTeSR Plus medium (STEMCELL Technologies). The iPSCs were differentiated into mesoderm-derived vascular smooth muscle cells (SMCs) over a 24 days differentiation protocol as previously described ^23^. Briefly, iPSCs (Day 0) were dissociated into single cells, seeded on Matrigel-coated dish cultured in E8BAC medium (E5 medium plus 5 ng/ml BMP4, 25 ng/ml Activin A, 19.4 mg/l insulin, 10 µM Y27632 and 1 µM CHIR99021) for 36 hours. Cells were dissociated again, seeded at low density on Matrigel and in grown in E6T medium (E5 medium supplemented with 19.4 mg/l insulin, 1.7 ng/ml TGF-β1 and 10 µM Y27632) for 18 hours (Day 3). Medium was then replaced with E5F medium (E5 medium supplemented with 19.4 mg/l insulin and 100 ng/ml FGF2) until day 8. From day 8 to day 11, the cells were treated with FVR medium (E5 medium supplemented with 19.4 mg/l insulin, 50 ng/ml VEGF and 5 µM RESV). Cells were then treated with E6-R medium (E5 medium supplemented with 19.4 mg/l insulin, 5 µM RESV, 25 µM Repsox) for 12 days, with a dissociation on day 16. After reaching the 24-day mark, iPSC-derived SMCs were maintained in E6-R medium, all of which were sourced from Thermo Fisher Scientific located in Waltham, MA, USA. Expression of SMC markers was verified using qPCR.

### Data visualization and statistical analyses

Unless otherwise noted, all statistical analyses and figures were performed in R (v4.2.2), using following packages: ggplot2 (v3.4.4), rtracklayer (v1.58.0), ggrepel (v0.9.3), data.table (v1.14.8), dplyr (v1.1.3), tidyr (v1.3.0). For TFBS prediction, a 51 bp sequence centered on rs13128814 was used as input to PERFECTOS-APE webserver (opera.autosome.org/perfectosape/scan), using HOCOMOCO-in vivo (v12) collection of position weight matrices as reference database^24^.

### Dual-luciferase enhancer reporter assay

To assess promoter activity, dual-luciferase reporter assays were conducted. Using BglII/HindIII restriction enzymes, the TK-minimal promoter was removed from pRL-TK (Promega, Madison, Wisconsin, USA) and inserted into the appropriate locations of pGL4.12 (Promega, Madison, Wisconsin, USA) to create the pGL-TK minimal promoter reporter plasmid. Between the Nhe1 and Xho1 sites of pGL-TK, a 947 bp DNA fragment including rs13128814 was amplified from the mix of human genomic DNA (primers: F: gctcgctagcACCTTTAAGTCTCGGCCTCC, R: tatcctcgagTCCCGGGTTCAAGCTATTCT). To generate rs13128814-A pGL4-TK plasmids, fusion PCR was performed with primers covering the variant (F: GGGCAGAGAACAGAGCCAAGTTCCTGTTTTGCTGC, R: GCAGCAAAACAGGAACTTGGCTCTGTTCTCTGCCC). A7r5 were seeded in 96-well plates with a concentration of 10,000 cells per well, after 24h, transfected with luciferase reporter constructs or containing the relevant promoter regions using FuGENE HD Transfection Reagent Complex (Promega, Madison, Wisconsin, USA). pCMV-NFIA1.1 (a kind gift from Richard Gronostajski, Addgene plasmid # 112698^25^) was transfected with luciferase-expressing plasmids at the same time for NFIA overexpression condition. 48h after transfection, the Firefly and Renilla luciferase were measured by performing Dual-Luciferase Reporter Assay System (Promega, Madison, Wisconsin, USA), following the instructions of the manufacturer. Signal was recorded using Mithras LB 940 Multimode Microplate Reader machine (Berthold Technologies, Bad Wildbad, DE). The enhancer capacity was calculated by the ratio of Firefly to Renilla luciferase. Statistical significance was evaluated using R studio t-test with unequal variances.

### siRNA knockdown

In 6-well plates, iPSC-derived SMCs (SKiPS-31.3 and 12.10 lines) and BJ fibroblast cells were seed at a density of 100,000 cells per well to reach a roughly 70–80% confluence for the siRNA knockdown tests. Silencer® Select N°1 negative control siRNA or predesigned Silencer® Select siRNAs for ZNF827 (s45694, s45695, s45696) (Thermo Fisher Scientific, Waltham, MA, USA) were used for the transfection process. For the 72-hour transfection process, Opti-MEM and Lipofectamine RNAiMAX transfection reagent (Invitrogen, Waltham, Massachusetts, USA) were used.

### Quantitative RT-PCR Analysis

Cells were collected and given a PBS rinse before being subjected to the quantitative RT-PCR analysis. RNA purification was carried out following the manufacturer’s instructions, utilizing the RNeasy Plus Mini kit (Qiagen, Hilden, Germany). Reverse transcription was carried out using the iScript cDNA Synthesis Kit (BIO-RAD, Hercules, California, USA). The final cDNA was used as a template for quantitative real-time PCR with the GoTaq qPCR master mix (SYBR Green), purchased from Promega in Madison, Wisconsin, USA. The PCR reactions were carried out on an Applied Biosystems StepOne Plus device, and primers were designed to detect *ZNF827* mRNA level produced using the internet tool https://primer3.ut.ee/ (F: TTTGAGTGTGATGTGTGCCA, R: ACCACTGTCCTGAGTTTCCT). The comparative CT approach in StepOne software was utilized to ascertain the relative amounts of gene expression. The housekeeping genes *GAPDH*, *ACTB*, and *SDHA* expression levels were used to standardize the results. To ensure accuracy, each reaction was carried out in triplicate. A two-sample t-test with unequal variances was used to determine the statistical significance of each result.

### RNA-Seq

PolyA+ mRNA libraries were prepared from 100ng-1µg of total RNA using QIAseq Stranded RNA library preparation kit (Qiagen) according to manufacturer’s instruction. Libraries were sequenced on a NextSeq500 instrument (Illumina, San Diego, CA, USA) using NextSeq 500/550 High Output Kit v2 (75 cycles). Reads were demultiplexed using bcl2fastq2 (v2.18.12) and adapters were trimmed using Cutadapt (v1.15). Reads were then mapped to human genome (GRCh38.104) using STAR aligner (v2.7.9a) with following options: --outFilterType BySJout --outFilterMultimapNmax 20 --alignSJoverhangMin 8 --alignSJDBoverhangMin 1 -- outFilterMismatchNmax 999 --outFilterMismatchNoverReadLmax 0.04 --alignIntronMin 20 - -alignIntronMax 1000000 --alignMatesGapMax 1000000. We used per gene read counts as direct input for differential expression analysis using DESeq2 (v1.38.3) package in R (v4.2.2), keeping only genes with mean read count over 1, using cell type (SMC or Fibroblasts) as a covariate^26^. We transformed the count matrix using variance stabilizing transformation with option blind=FALSE. Differentially expressed genes were determined using res function and log2 fold changes were determined using lfcShrink function with ashr shrinkage estimator ^27^. Gene ontology term enrichments were calculated using clusterProfiler package (v4.6.2).

### Cell viability assay

Cells (5,000 cells/well) were seeded in 96-immuno white immune plates (Thermo Fisher Scientific, Waltham, MA, USA) with 16 replicates. CellTiter-Glo 2.0 assay (Promega, Madison, Wisconsin, USA) was used to measure cell viability. Plates were loaded with CellTiter-Glo reagents and the luminescent signal was recorded using Mithras LB 940 Multimode Microplate Reader machine. Measurements of luciferase were performed 4h, 24h, 48h, 72h and 144h after transfection. The viability of all luciferase measurements was normalized to the values at 4 h. Statistical significance was evaluated using Student’s two sample t-test with unequal variances.

## Results

### A single variant may cause association to multiple cardiovascular traits and disease at *ZNF827* locus

In our previous study, the *ZNF827 locus* was found to be associated with spontaneous coronary artery dissection (SCAD), with rs1507928 as lead variant. To identify whether other traits or diseases could potentially be associated to the same genetic variants, we looked up GWAS catalog for associations involving variants in high linkage disequilibrium with SCAD lead variant rs1507928 (r^2^ > 0.7 in European population of 1000G reference panel, **Supplementary Table 1**). We found several cardiovascular traits and diseases including coronary artery disease (CAD, lead SNP rs10006310, *P* = 3×10^-9^)^28^, systolic blood pressure (SBP, lead SNP rs4835266, *P* = 8×10^-17^) ^29^ and ascending aorta diameter (AAdia, lead SNP rs11100902, *P* = 6×10^-9^)^16^ (**Supplementary Table 2**). To determine whether these different associations may be caused by a same variant, we retrieved summary statistics of genetic association to 7 cardiovascular traits/disease with significant association. Each association involves multiple non-coding genetic variants in high linkage disequilibrium associated with these traits and diseases at *ZNF827* locus (**Figure 1A, Supplementary Figure 1**). Using HyPrColoc multi-trait colocalization, we found these 7 traits to cluster together with an overall probability that they share a single causal variant estimated to 85% (**Figure 1B**). Under this hypothesis, a single variant (rs13128814) appeared as the most likely to be causal for all 7 associations (PP_rs13128814_ = 0.91, **Figure 1B**). This variant is significantly associated to SCAD, SBP, PP, and various aorta measurements, while it was also suggestively associated to CAD, fibromuscular dysplasia and diastolic blood pressure (**Figure 1C**). Interestingly, rs13128814 risk allele for SCAD (rs13128814-A) appeared as a protective allele for CAD and was associated to lower blood pressure, while it was associated to increased size of the aorta (**Figure 1C, Supplementary Table 3**). rs13128814 is located within the 4^th^ intron of ZNF827 gene and overlapped histone marks characteristics of active regulatory elements in coronary artery tissue (**Figure 1D**). Using single-nucleus chromatin accessibility map from diseased and healthy coronaries^20^ **(Supplementary Figure 2)**, we observed that this regulatory element seems particularly active in smooth muscle cells and fibroblast cells types, while it was not accessible in endothelial or immune cells (**Figure 1D**).

**Figure 1.**
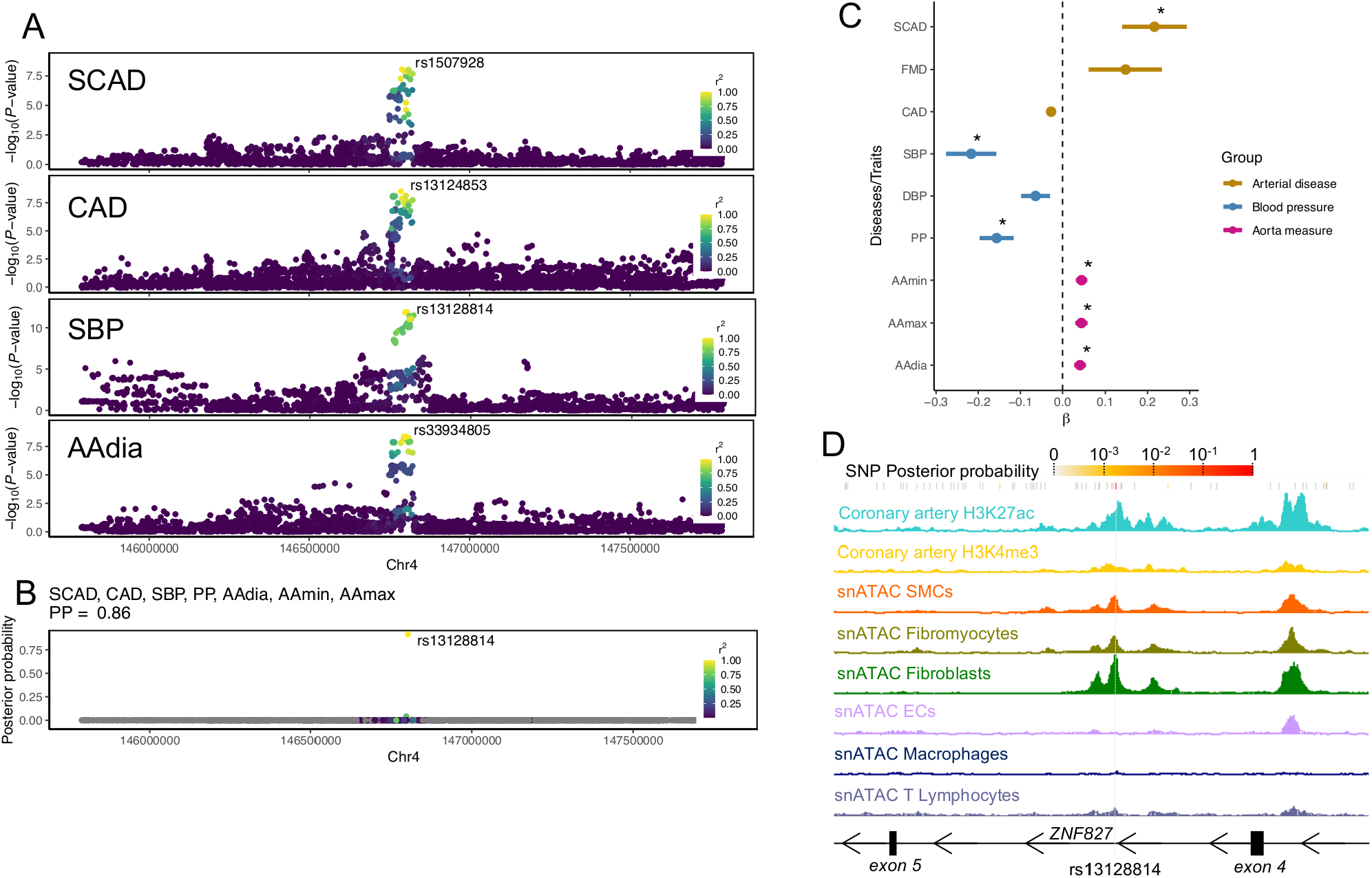
Genetic associations and functional annotation at ZNF827 locus. **A**. Local genomic plots represented the genetic association of common variants in *ZNF827* locus to spontaneous coronary artery dissection (SCAD), coronary artery disease (CAD), systolic blood pressure (SBP), and ascending aorta diameter (AAdia). Top variant is indicated for each association, and color indicates the linkage disequilibrium (r2 in European population of 1000 Genomes reference panel) of all variants to top variant. **B**. Bayesian posterior probability for each variant to be causative of the genetic associations with SCAD, CAD, SBP, pulse pressure (PP), AAdia, ascending aorta minimal maximal areas (AAmin, AAmax). **C**. Forest plot representing beta and 95% confidence interval of the associations between rs13128814 (A allele) and 9 cardiovascular diseases/traits. FMD: Fibromuscular dysplasia. DBP: diastolic blood pressure. A star indicates P-value of association lower than 5×10^-8^. **D**. Genome browser visualization of histone chromatin-immunoprecipitation and single nuclei snATAC-Seq read densities in the region surrounding rs13128814. The dashed grey line highlights rs13128814 exact position.

### rs13128814 risk allele is required for enhancer activity

To determine whether enhancer activity may be affected by rs13128814 genotype, we cloned a 948 bp genomic region centered on the observed open chromatin region in a luciferase reporter plasmid. This region overlapped 3 common genetic variants, all significantly associated with SCAD and in high LD in European population, rs13128814, rs17020769 and rs1979974 (**Figure 2A**). First, we compared the transcriptional activity of regions comprising all protective allele and all risk allele. We observed that the region containing all risk allele was associated to an increased transcriptional activity compared to an empty vector, while the region with protective allele had no transcriptional activity (**Figure 2B**). Then, we compared the activity of the region with all protective alleles with the same region with only rs13128814 risk allele (A). We observed that this allele alone was sufficient to increase transcriptional activity to the same level as with all risk alleles (**Figure 2B**), suggesting that rs13128814-A allele is required to have active transcriptional activity in this enhancer region.

**Figure 2.**
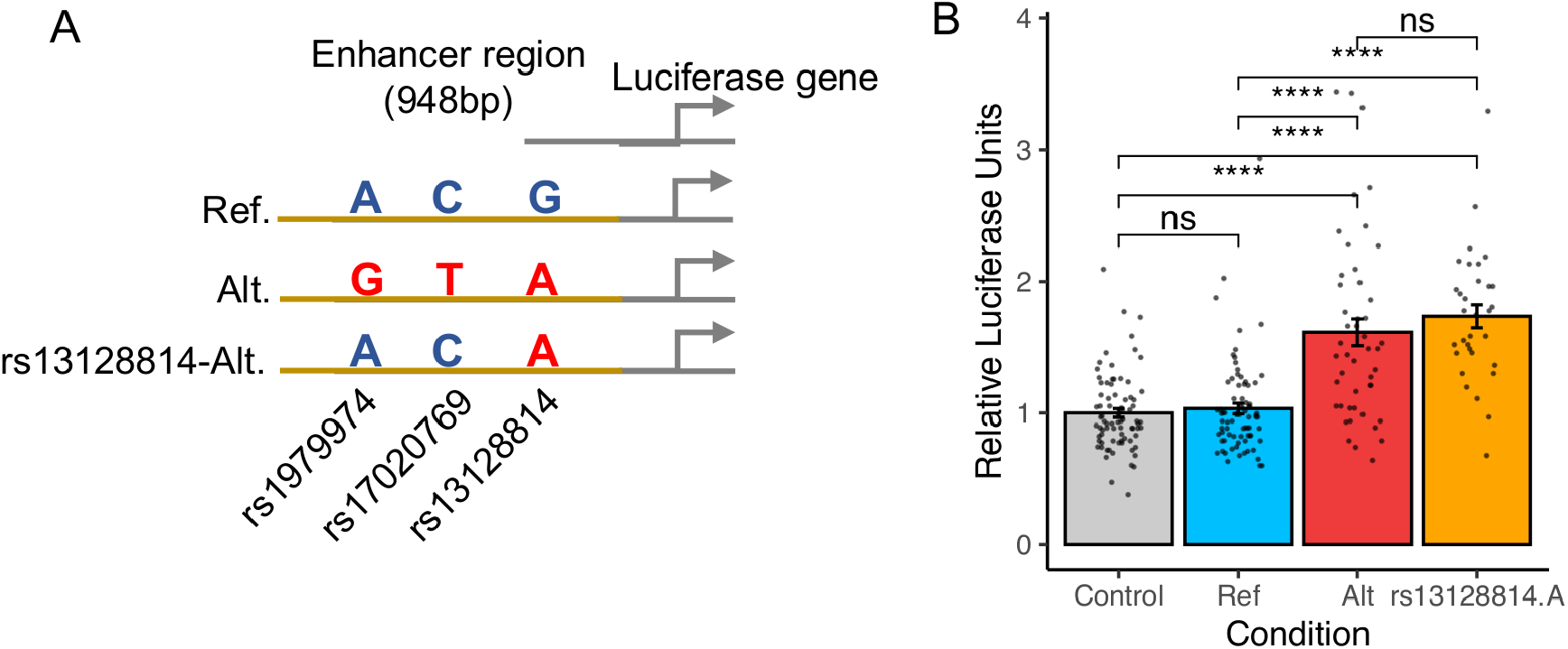
Transcriptional activity of rs13128814-surrounding region. **A.** Illustration of the region encompassing rs13128814 cloned in a luciferase reporter plasmid. rsID and genotype of the 3 common variants overlapping this DNA fragment are indicated for each construct. **B:** Luciferase activity measured though overexpression of indicated constructs in A7r5 rat SMCs. Wilcoxon Rank Sum test (two-tailed): ****: *P*<10^-4^; ns: *P*≥0.05.

### rs13128814 risk allele creates a binding site NF-1 transcription factors

We used PERFECTOS-APE algorithm to predict transcription factor binding sites (TFBS) potentially affected by rs13128814 genotype, using in vivo sites from HOCOMOCO v12 as candidate TFBS^24^. We found 12 candidate TFBS, corresponding to 10 candidate TF, predicted to recognize differentially the sequence including rs13128814 (log2 Fold Change > 4, **Figure 3A, Supplementary Table 4**). The first 2 candidate TFBS corresponded to sites of Nuclear Factor 1 family (NF-1) NFIA and NFIC, with other differential TFBS predicted for other NF-1 factors NFIB and NFIX. rs13128814-A allele perfectly matched NF-1 binding motif while rs13128814-G disrupted this motif (**Figure 3B**). NF-1 family factors were highly expressed in coronary artery tissues from GTEx database, while other candidate TF were poorly expressed in this tissue (**Figure 3C**), and particularly present in SMC and Fibroblast cell types (**Supplementary Figure 3**). To determine whether NF-1 factors could activate rs13128814-related enhancer, we overexpressed mouse NFIA in the previously described luciferase assay (**Figure 3D**). We observed that NFIA overexpression increased luciferase expression in absence of any regulatory region, suggesting NFIA may bind sites in the luciferase plasmid (**Figure 3D**). We however observed an increased activation of transcription when NFIA was overexpressed with the plasmid containing rs13128814-A allele, but not with the plasmid containing rs13128814-G allele (**Figure 3D**).

**Figure 3.**
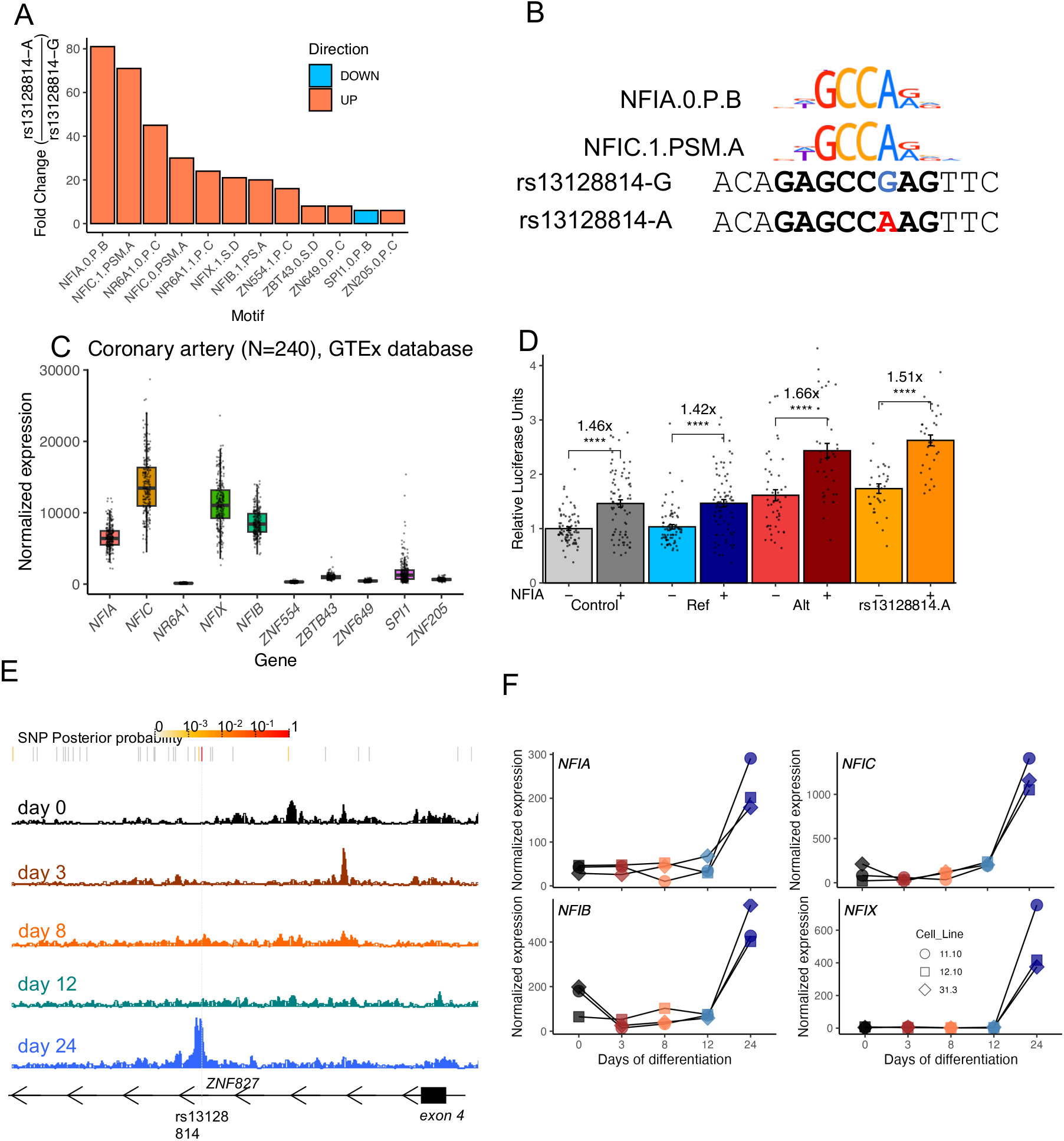
rs13128814 disrupts a potential binding site for NF-1 transcription factors. **A.** Candidate TFBS affected by rs13128814. Barplot represents the ratio of P-value of similarity of DNA sequence containing rs13128814-A allele to each TFBS with its counterpart with G-allele. Color indicates if the similarity to TFBS is higher (red) or lower (blue) with rs13128814-A allele. Approximate P-values for position weight matrices were estimated using threshold method as implemented in PERFECTOS-APE package^48^. **B.** Alignment of DNA sequence close to rs13128814 with NFIA and NFIC binding motifs from HOCOMOCO v12 (“in vivo” TFBS). **C.** Normalized expression of candidate transcription factors in 240 coronary artery samples from GTEx database. Boxes represents the 25^th^,50^th^ and 75^th^ centiles of the distribution. Lower and upper whiskers indicate a maximal distance of 1.5 times the interquartile range from first and third quartile, respectively. **D.** Luciferase activity measured though overexpression of indicated constructs in A7r5 rat SMCs, with or without pCMV-Nfia. Fold-change is indicated over the graph. Wilcoxon Rank Sum test (two-tailed): ****: *P*<10^-4^. **E.** Genome browser visualization of chromatin accessibility in iPSCs (day 0) and along differentiation to SMCs (days 3 to 24). **F.** Normalized expression of genes coding for Nuclear factor-1 transcription factors in iPSCs (day 0) and along differentiation to SMCs (days 3 to 24).

We recently observed that differentiation of vascular SMCs using RepSox, a TGF-β inhibitor, could lead to a regulatory phenotype closer to artery tissue. Indeed, we observed that rs13128814-associated regulatory element became accessible during the last step of this differentiation protocol, concomitantly with the increased expression of SMC markers (**Figure 3E, Supplementary Figure 4**). Interestingly, this activation was accompanied by an increased expression of NF-1 transcription factors, further supporting that these TF may be required for the activation of this regulatory element (**Figure 3F**).

### *ZNF827* acts as a major regulator of gene expression in SMCs and fibroblasts

To determine which could be the target gene at ZNF827 locus, we looked-up rs13128814 in GTEx database. We found that rs13128814-A allele was associated to higher expression of ZNF827 in tibial arteries and aorta, the two most powerful arterial datasets in GTEx (**Figure 4A-B).** No association was however observed in coronary arteries. We performed a colocalization of SCAD GWAS association with the eQTL association and we confirmed SCAD association colocalized with ZNF827 eQTL both in Tibial Artery (**Figure 4C**, PP.H4.abf = 0.96) and Aorta (PP.H4.abf = 0.94), supporting that these genetic associations may have a single common causal variant. The expression of other nearby genes was not associated to rs13128814 in any tissue.

**Figure 4.**
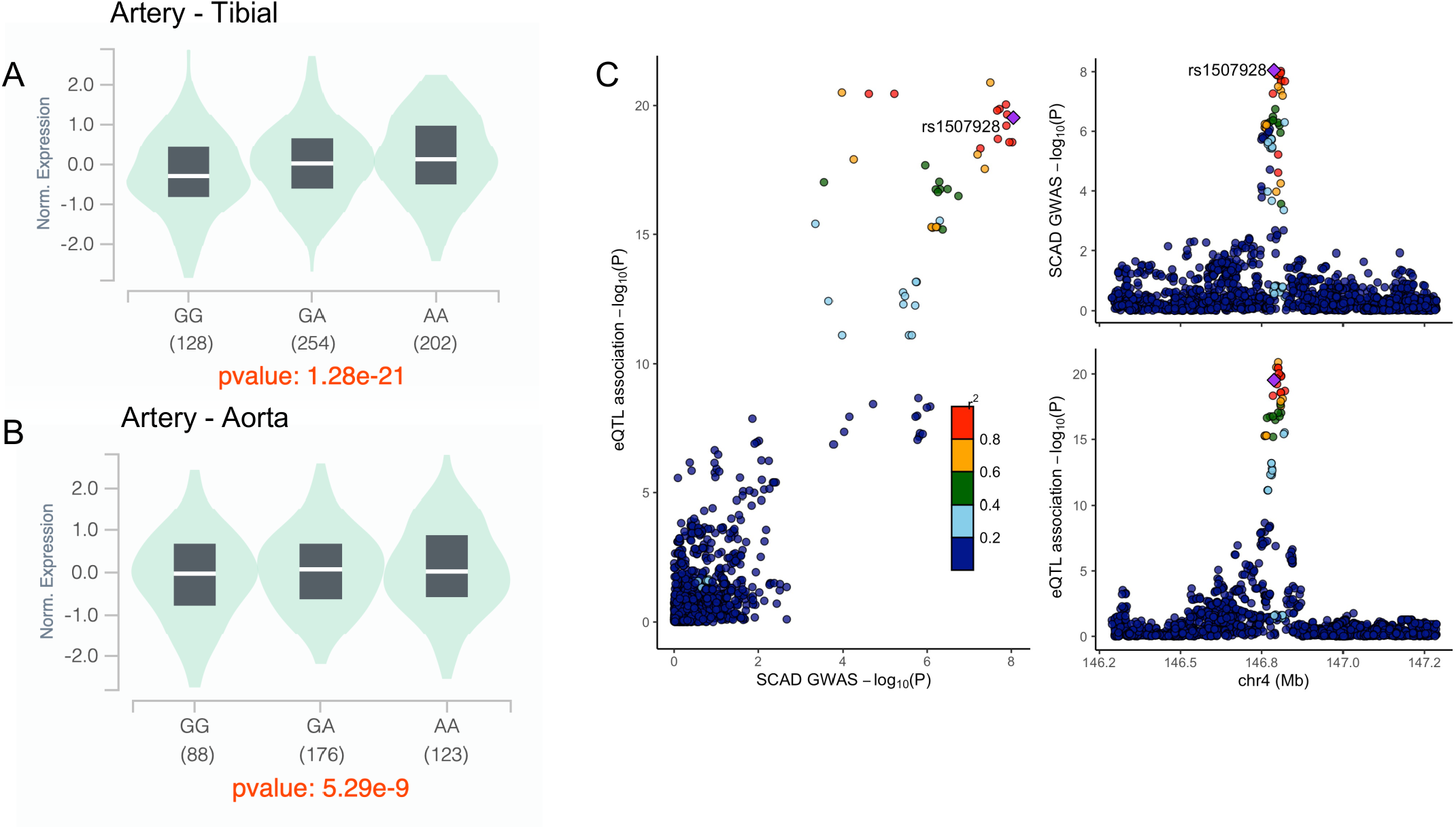
rs13128814 affects ZNF827 expression in artery tissues. **A-B.** Violin plots illustrating the association of rs13128814 genotype with *ZNF827* normalized expression in tibial artery tissue (**A**, *P*=1.3 × 10^-21^), and aorta (**B**, *P*=5.3 × 10^-9^). **C.** Colocalization of genetic association signals for SCAD and ZNF827 eQTL in Tibial artery. Approximate Bayes factor posterior probability for two association to share a single causal variant was estimated to 96%.

The function of *ZNF827* in arterial tissue is not known. To get insight into its potential role in cardiovascular disease, we knocked down *ZNF827* expression in iPSC-derived SMCs (2 clones, 3 replicates) and BJ primary fibroblasts (3 replicates) using siRNAs. We collected RNA 48h post-transfection, confirmed *ZNF827* knockdown using qPCR (**Supplementary Figure 5**), and analysed gene expression using high-throughput sequencing. We identified 437 differentially expressed genes including both SMCs and fibroblasts (*P.adj* < 0.05) with a majority of genes (270) downregulated following *ZNF827* knockdown (**Figure 5A, Supplementary Table 5**). Differential expression was highly correlated between SMCs and fibroblasts (R = 0.36, *P* < 2×10^-16^, **Figure 5B**). Pathway enrichment analysis showed an overrepresentation of genes involved in regulation of macroautophagy, with key autophagy factors such as *ATG5* (*Autophagy protein 5*) and *NBR1* (*Next to BRCA1 gene 1*) amongst most downregulated genes (**Supplementary Table 6, Figure 5C-D**). Beyond enriched pathways, several of the most dysregulated genes following *ZNF827* knockdown play important roles in atherosclerosis and arterial disease including *Caveolin-1* (*CAV1*)^30,31^, *Osteoprotegerin (TNFRSF11B)*^32,33^, *Cytoskeleton-associated protein 4 (CKAP4)*^34^ and *Transcription factor E2F4 (E2F4)*^35^.

**Figure 5.**
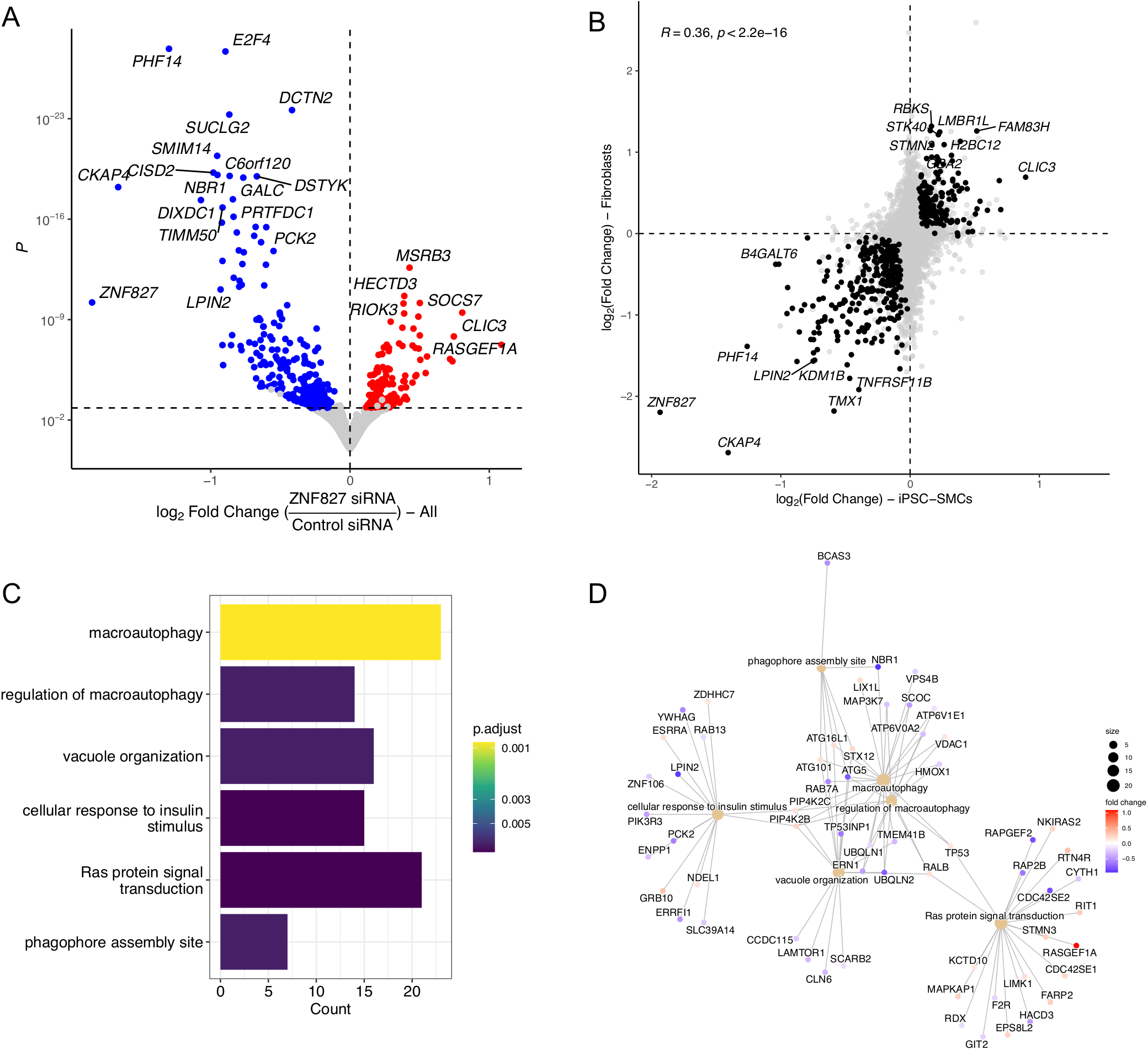
ZNF827 regulates multiple genes in SMCs and fibroblasts. **A.** Volcano plot representation of differential expression following ZNF827 knockdown in iPSC-derived SMCs (2 clones, 3 replicates each) and BJ Fibroblasts (3 replicates). Log2 Fold Change is represented on *x* axis, while P-value is represented on *y* axis (log scale). Differentially expressed genes (*P.adj* < 0.05) are highlighted in blue (downregulated genes) or red (upregulated genes). **B.** Scatterplot representation of differential expression in iPSC-derived SMCs (*x* axis) versus BJ Fibroblasts (*y* axis). Consistent differentially expressed genes are highlighted in black. **C.** Top enriched gene ontology pathways among differentially expressed genes. **D.** Network representation of differentially expressed genes involved in enriched pathways.

Although differential expression was highly correlated between SMCs and fibroblasts, a large group of genes was specifically regulated only in fibroblasts (**Supplementary Figure 6**). Pathway analyses showed an important enrichment of downregulated genes involved in mitosis and cell cycle regulation, whereas upregulated genes were enriched in pathways related to protein synthesis (**Supplementary Figure 7**). Indeed, reduced cellular proliferation was observed in BJ fibroblasts following *ZNF827* knockdown (**Supplementary Figure 8)**.

## Discussion

In this study, we focused on the exploration of regulatory mechanisms in a genetic risk locus recently associated to SCAD, CAD, blood pressure and aorta size. First, we demonstrated that the genetic association signals are consistent with the hypothesis that a single variant could cause the association to all these vascular traits and diseases. We identified rs13128814, a variant located in a regulatory element active in coronary artery SMCs and fibroblasts, as the most likely causal variant. We showed that rs13128814 risk allele was associated to higher transcriptional activity and higher sensitivity to overexpression of NF-1 transcription factor *NFIA*, and increased expression of *ZNF827* gene. We observed that *ZNF827* knockdown caused major transcriptomic changes, including several genes involved in atherosclerosis and vascular remodeling. Altogether, our study provides a plausible mechanism to understand the pleiotropic genetic association at *ZNF827* locus.

Among key findings in this study, we found that rs13128814 SCAD risk allele, which is the protective allele for CAD, creates a consensus binding site for transcription factors of the NF-1 family. Interestingly, the role of this family of transcription factors in vascular biology has not been studied in detail to our knowledge. Deletion of NF-1 transcription factors in mice was associated to a wide variety of phenotypes, but no cardiovascular alterations were reported to date^36–39^ However, NF-1 binding sites are amongst the most strongly enriched in active regulatory regions of vascular SMCs and endothelial cell^20^. NF-1 factors *NFIA* and *NFIB* were also recently identified as key driver genes in female specific gene regulatory networks in atherosclerosis, mostly through SMC related pathways^40^. The large compensation between NF-1 factors, which all recognize the same DNA sequence, and are expressed at high levels in vascular SMCs, could explain the lack of cardiovascular phenotypes related to inactivation of individual NF-1 factors. Nevertheless, these recent results and our study all point to the fact that NF-1 factors could play a key role in pathological vascular remodeling.

We identified *ZNF827* as the most likely target gene at this locus. *ZNF827* is expressed in all tissues and predicted to encode a DNA-binding protein, but its potential function in the vasculature is not known. Several studies have found that ZNF827 could regulate various processes such as epithelial-to-mesenchymal transition^10^, splicing^10^, alternative lengthening of telomeres^41^ through homologous recombination^42^ and neuronal differentiation^12^. Studies also found that the function of *ZNF827* gene could be mediated both by its protein product and by circular RNA^12^. Here, through a knockdown approach we found that *ZNF827* downregulation induced a major transcriptional response with over 900 genes dysregulated in iPSC-derived SMCs and more than 3000 dysregulated genes in fibroblasts. The difference between fibroblasts and SMCs might be due to a large effect of *ZNF827* knockdown to slow down cell cycle, whereas iPSC-derived SMCs proliferate at a low pace^21^.

Among genes commonly regulated in iPSC-derived SMCs and fibroblasts, we found several genes particularly relevant to vascular disease. *TNFRSF11B*, which encodes osteoprotegerin, was amongst the most downregulated genes both in SMCs and fibroblasts. This gene was identified as one of the most specific markers of fibromyocytes, a subtype of SMCs thought to play a key role in atherosclerosis^32^. Osteoprotegerin was also found to act as an antagonist of vascular calcification, although its role is complex and involves interaction between vascular SMCs, endothelial cells and immune system^33,43^. The most downregulated gene following *ZNF827* knockdown is *CKAP4*, which encodes the Cytoskeleton-associated protein 4. CKAP4, also known as cytoplasmic protein 63, is a transmembrane and endoplasmic reticulum protein acting as a receptor to several proteins such as tissue plasminogen activator^44^, Wnt inhibitor Dickkopf-1^45^ or nexin 17^46^, with major implications in angiogenesis and SMC function. *CKAP4* was recently found to regulate vascular SMC survival in the context of abdominal aortic aneurysm ^34^. Several other of top *ZNF827* target genes play important roles in vascular biology, such as *CAV1* or *E2F4*^30,31,35^.

Our study suffers from several limitations. First, we could not demonstrate that NF-1 binding is allele specific *in vivo*, or that NF-1 directly regulated the expression of *ZNF827*. It is thus possible that the binding of other transcription factors may be affected by rs13128814 genotype. Second, we used a knockdown approach to identify the target genes of *ZNF827*, but we could not evaluate the long-term effects of *ZNF827* depletion or overexpression. We attempted to create knockout cell lines in iPSCs using CRISPR-Cas9, but despite several attempts we could never obtain mutant clones. One possibility is that *ZNF827* is essential to the survival of pluripotent cells. Indeed, based on GnomaD database (v4), *ZNF827* appears to be under strong selection pressure with very few deleterious mutations in the population (pLI = 1)^47^, suggesting it may be an essential gene. Similarly, we could not overexpress *ZNF827* without observing major cell death after a few days, limiting further explorations of its function. Finally, all our work was performed either from genetic data or from cell culture. Exploration of *ZNF827* in animal models or more complex cellular systems would be required to fully understand the role it may play in vascular remodeling and dissection.

Altogether, our work provides an exploration of regulatory mechanisms at a genetic locus involved in SCAD, CAD, blood pressure regulation and aorta dimensions. Several other genetic loci are associated to two or more of these cardiovascular traits and diseases, suggesting common biological pathways could be involved. Future studies focusing on these different loci may shed light on the shared genetic basis of cardiovascular diseases.

## Acknowledgements

We thank Alberto Tezza for its technical help in the preparation of this manuscript.

