## Supplemental Material for "*ZNF827* pleiotropic cardiovascular risk locus involves regulation by Nuclear factor-1"

**Supplementary Figure Legends**

**Supplementary Figure 1.**
Local genomic plots represented the genetic association of common variants in *ZNF827* locus to pulse pressure (PP), ascending aorta maximal area (AAmax) and ascending aorta minimal area (AAmin). Top variant is indicated for each association, and color indicates the linkage disequilibrium (r2 in European population of 1000 Genomes reference panel) of all variants to top variant.

**Supplementary Figure 2.**
**A**: UMAP plot of single-nucleus ATAC-seq. Clustering and annotation of single-nucleus was performed based on the pipeline and annotation strategy provided by authors[20]. B: Gene activity for marker genes associated to SMCs (*ACTA2, MYH11*) endothelial cells (*PECAM1*), fibroblasts (*LUM*), macrophages/monocytes (*CD14*) and T-cells (*RXRB*)

**Supplementary Figure 3**
Representative UMAP plots showing single cell populations profiled in single nuclei RNA-Seq analysis of diseased human coronary arteries [32] visualized using PlaqView[49], and featureplots of *ZNF827, NFIA, NFIB, NFIC* and *NFIX.*

**Supplementary Figure 4**
Normalized expression of pluripotent stem cell markers coding genes (*NANOG*, *SOX2*, and *POU5F1*), mesenchymal stem cell markers coding genes (*SNAI1*, and *PDGFRB*), vascular smooth muscle cell markers coding genes (*ACTA2*, *TAGLN*, *MYH11*, and *SMTN*) and *ZNF827* gene in iPSCs and along differentiation to SMCs in iPSCs (day 0) and along differentiation to SMCs (days 3 to 24).

**Supplementary Figure 5**
Relative expression of *ZNF827* in iPSCs-SMCs and fibroblasts under siRNA non-target control or *ZNF827* knockdown condition.

**Supplementary Figure 6**
**A-B:** Volcano plot representation of differential expression following *ZNF827* knockdown in iPSC-SMCs (**A**, 2 clones, 3 replicates each) and BJ Fibroblasts (**B**, 3 replicates). Log2 Fold Change is represented on *x* axis, while P-value is represented on *y* axis (log scale). Differentially expressed genes (*P.adj* < 0.05) are highlighted in blue (downregulated genes) or red (upregulated genes). **C:** Venn diagram illustrating the proportion of differentially expressed genes shared between SMCs and fibroblasts or restricted to one cell type

**Supplementary Figure 7**
Barplot representation of gene ontology pathways enriched amongst genes downregulated or upregulated following *ZNF827* knockdown in BJ fibroblasts. Top 20 pathways are represented. Bar color represent the adjusted P-value.

**Supplementary Figure 8**

Measurement of cell viability (mean±SEM,) of BJ fibroblasts treated with control siRNA (grey dots, 16 replicates) and two different *ZNF827* siRNAs (green squares and diamonds, 16 replicates each) at 4h, 24h, 48h, 72h and 144h. The unadjusted t-test was used at each time point: *p<0.05, **p<0.01, ***p<0.001, ns: not significant.

### Supplementary Figure 1

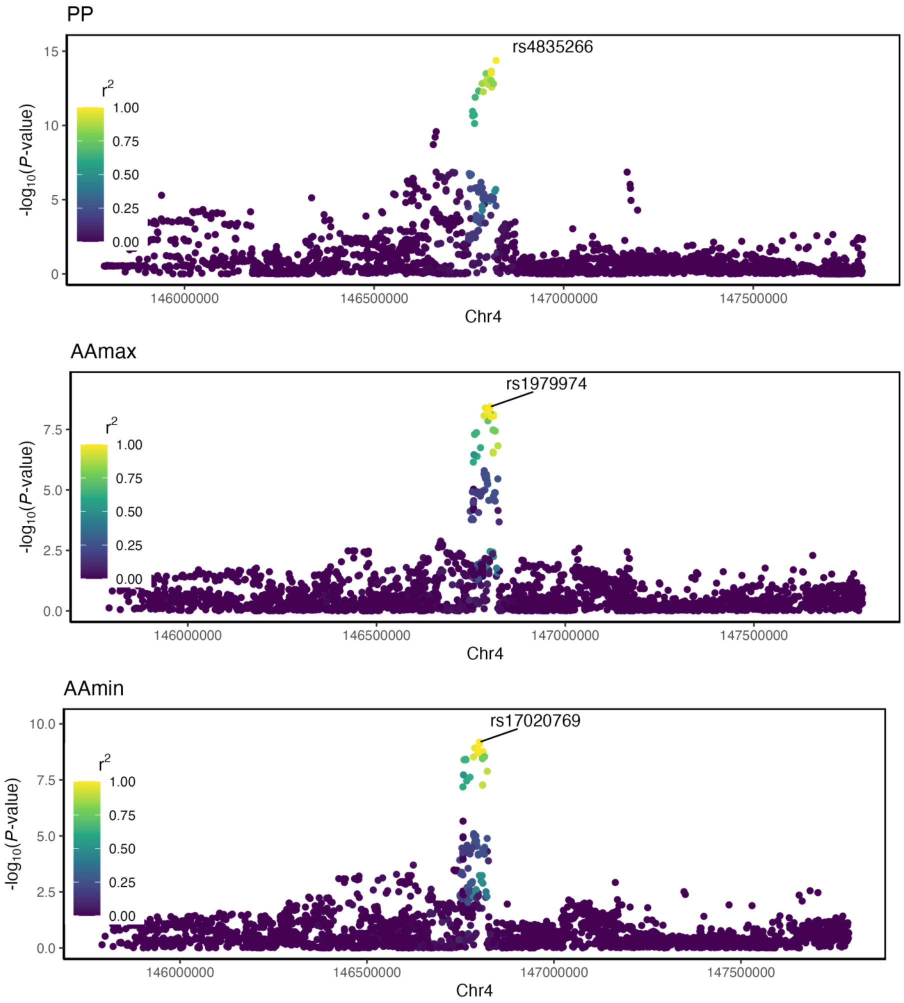

Supplementary Figure 2

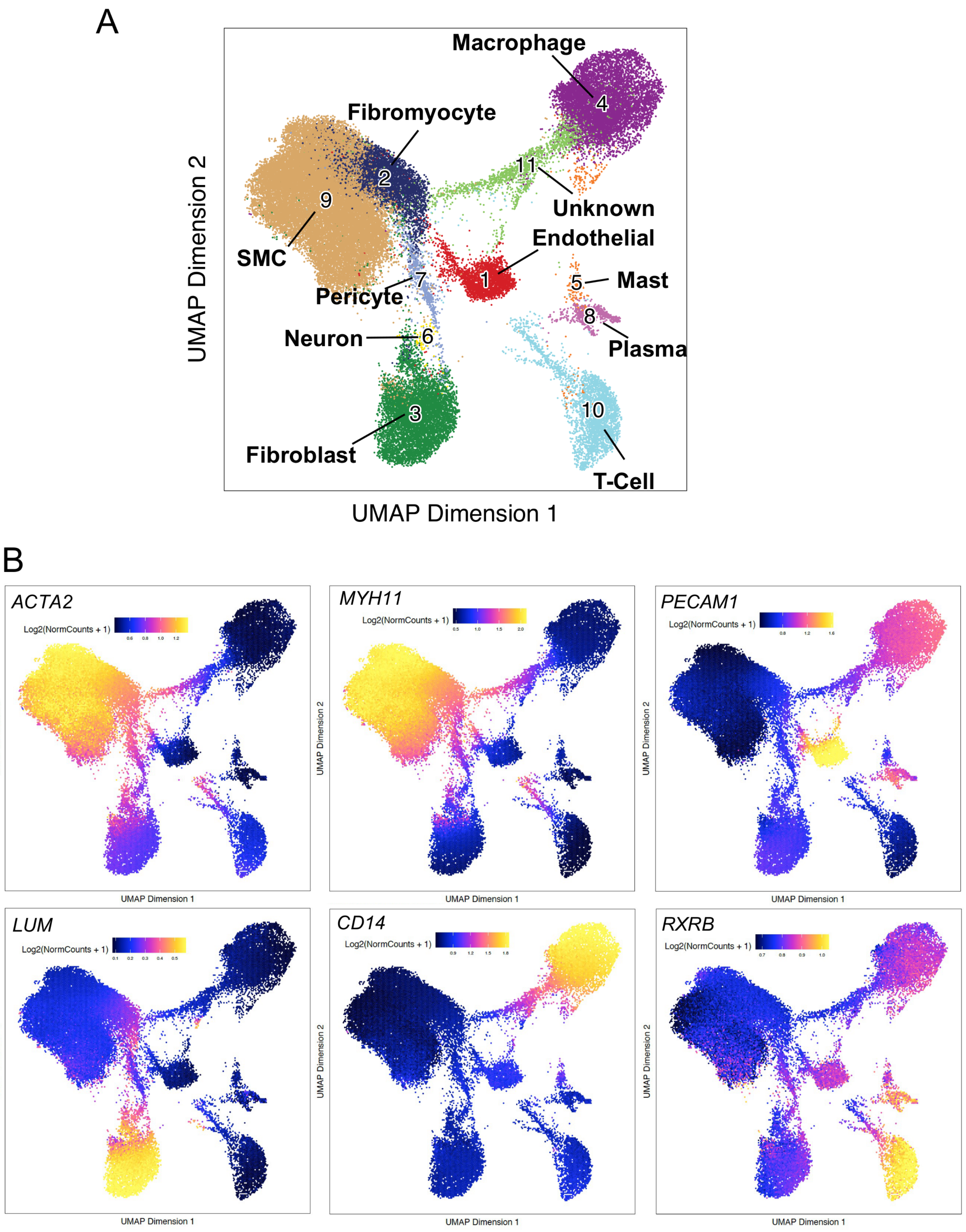

### Supplementary Figure 3

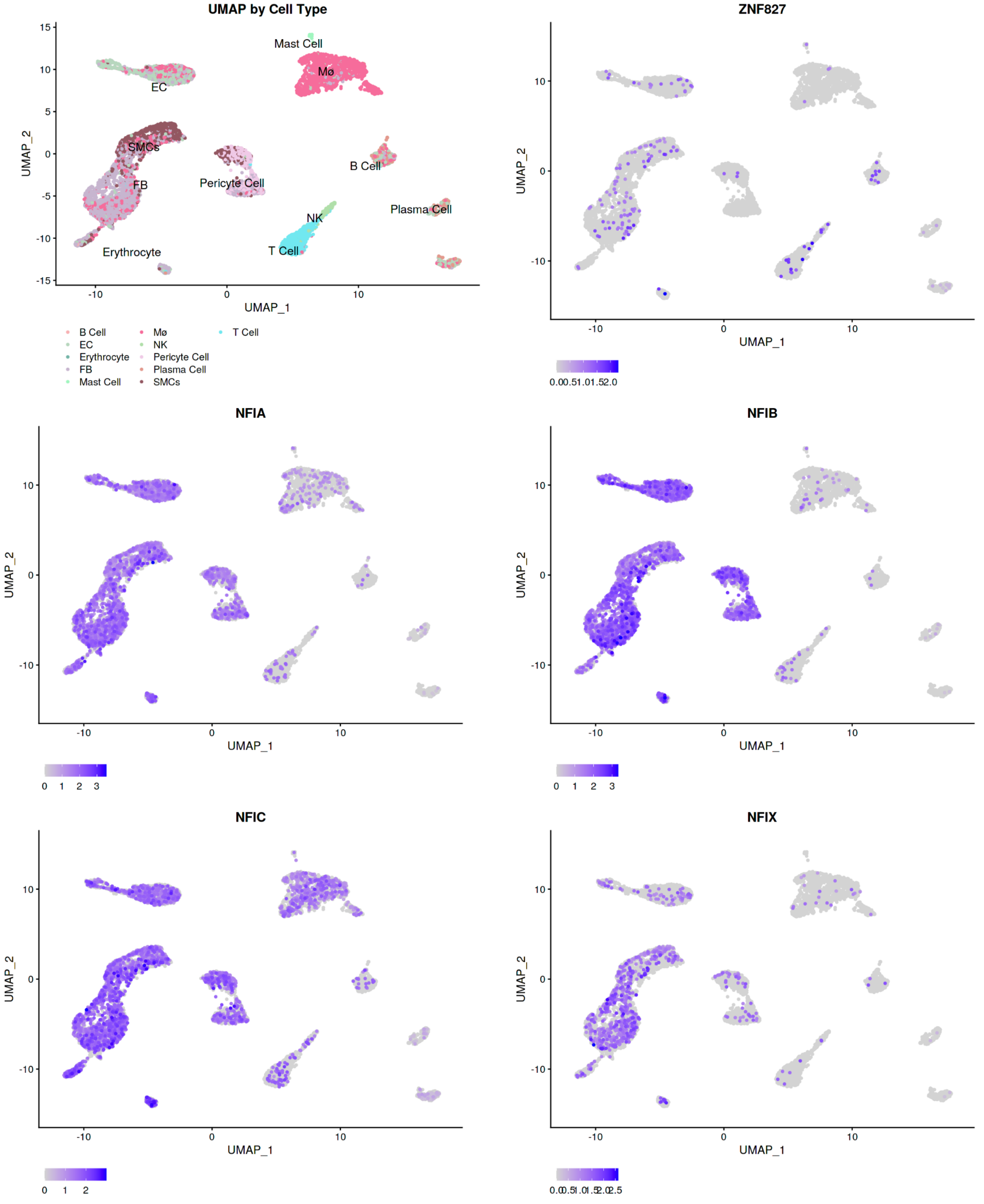

### Supplementary Figure 4

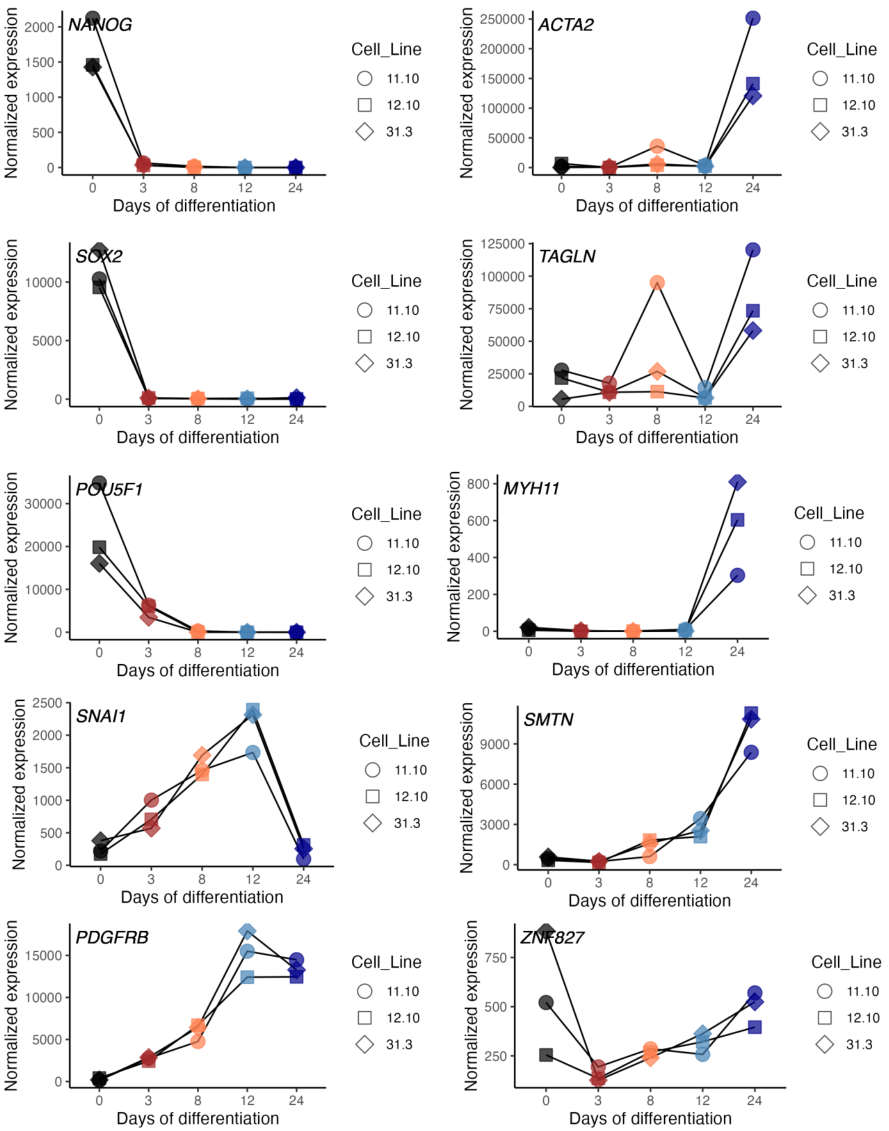

### Supplementary Figure 5

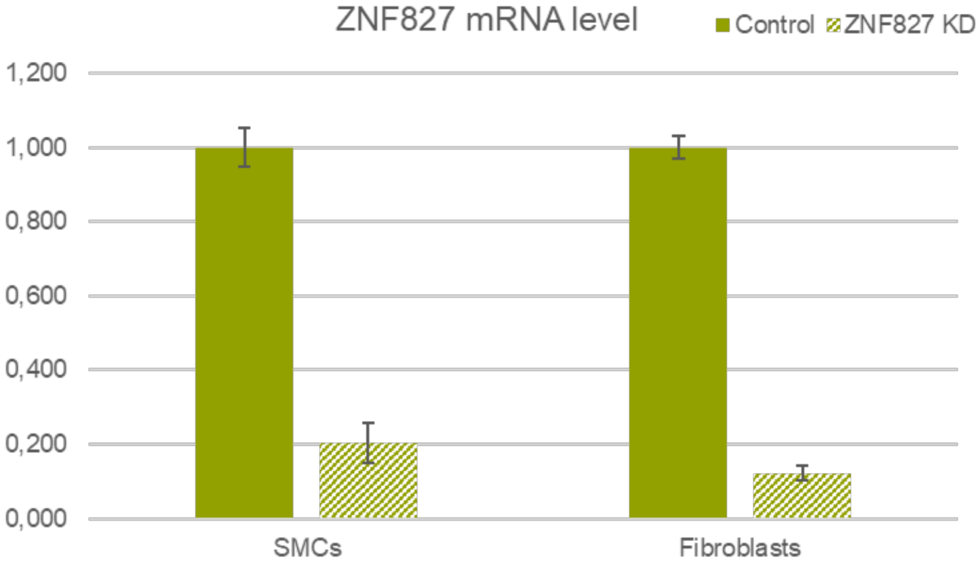

### Supplementary Figure 6

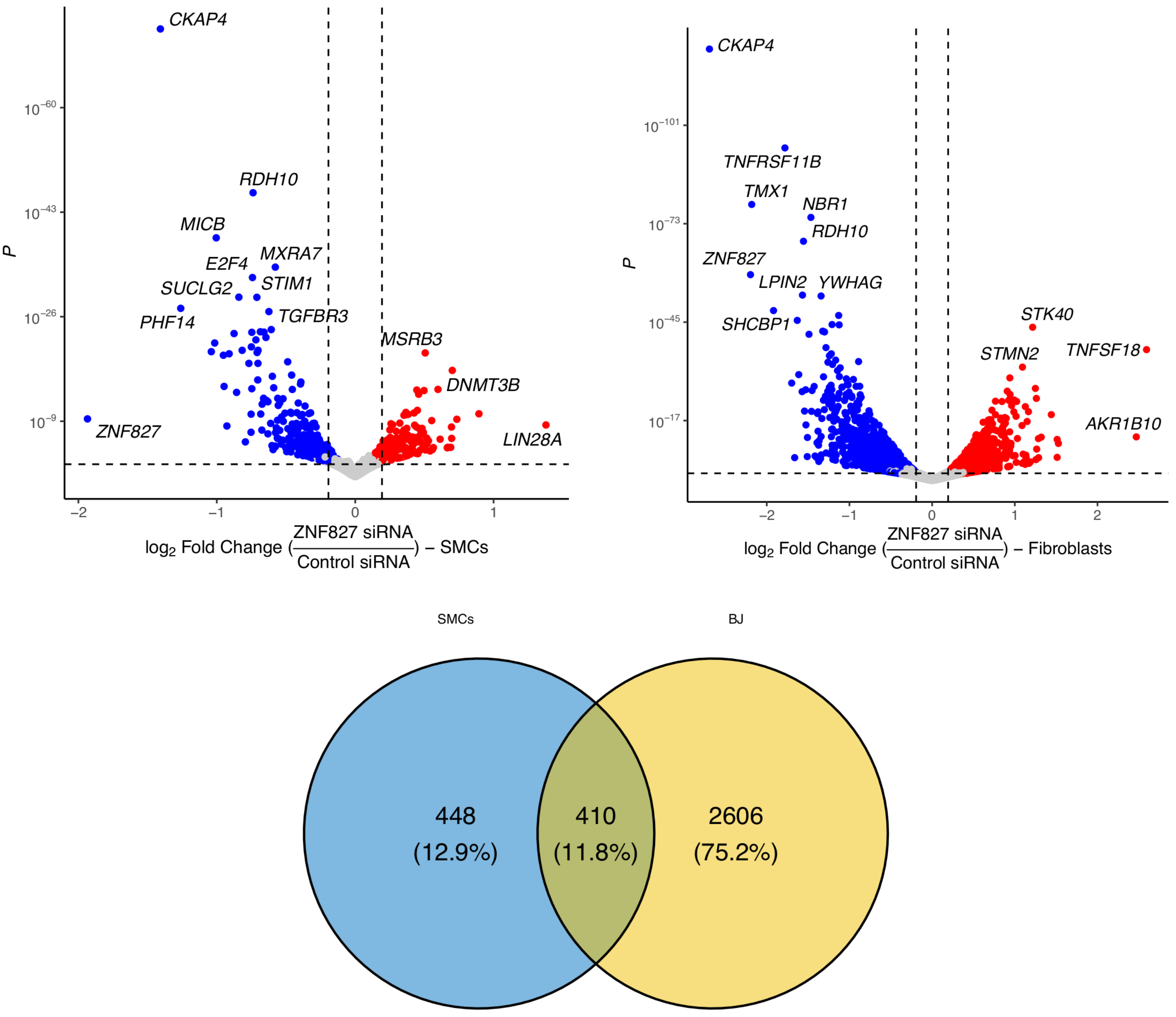

### Supplementary Figure 7

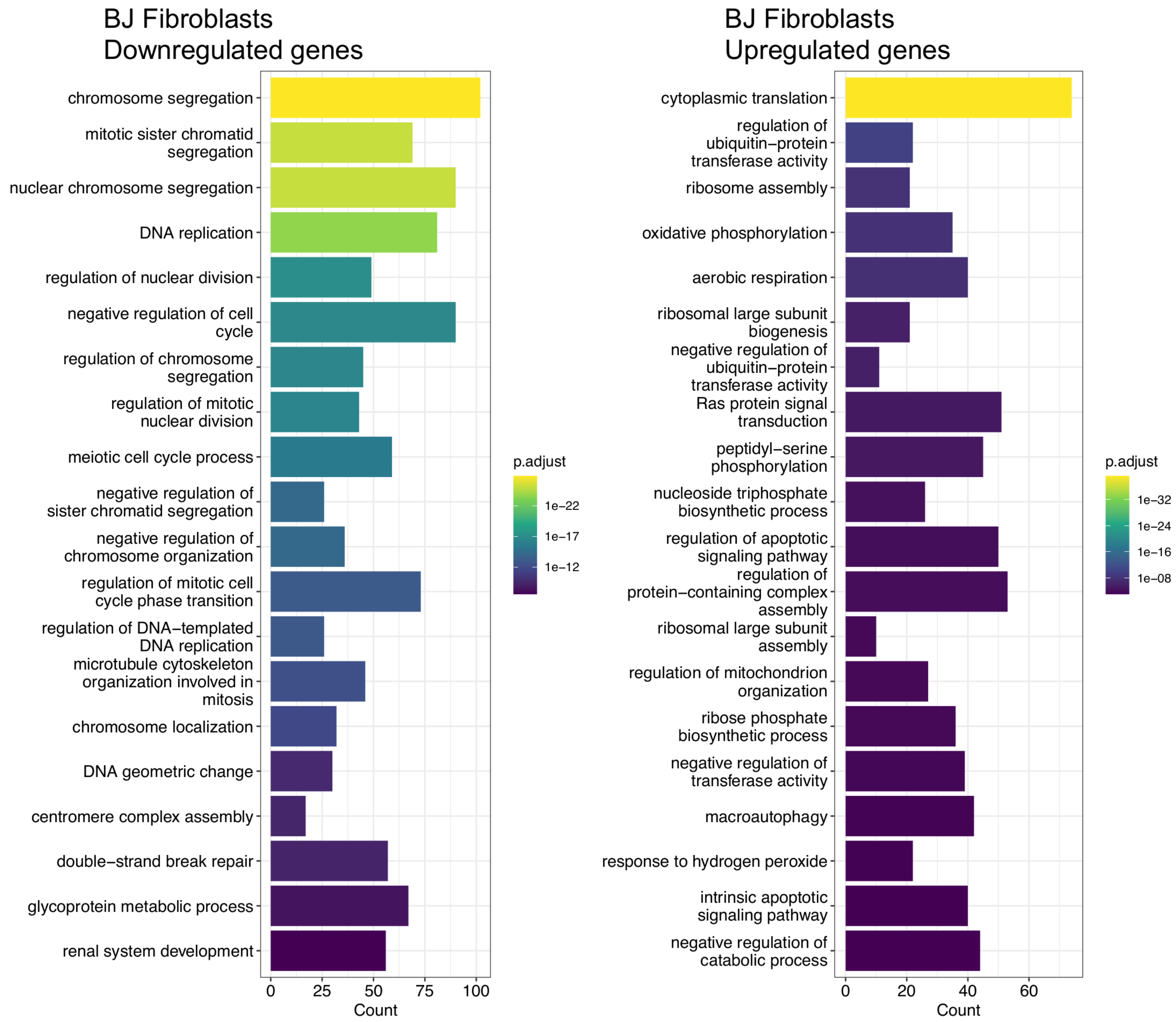

### Supplementary Figure 8

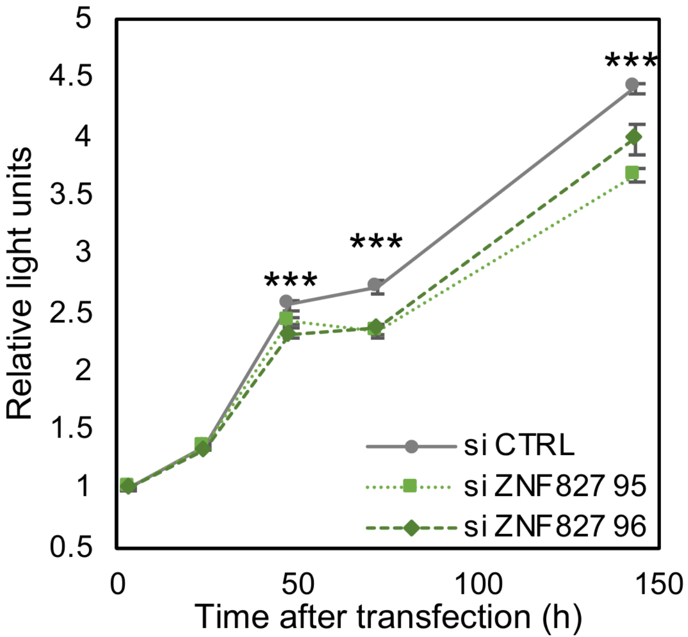

**Supplementary Tables Legends**

**Supplementary Table 1**
**SNP proxies at *ZNF827* locus.** All SNPs in high linkage disequilibrium ($r^2$>0.7) of SCAD lead SNP (rs1507928) in the European population of 1000Genomes reference panel are indicated. Proxies were retrieved using LDlink webserver.

**Supplementary Table 2**
**Lookup for associated traits in GWAS Catalog database.** GWAS Catalog was looked-up for proxies from Supplementary Table 1. For each trait/disease, the association with lowest P-value is shown, when the same association was found in several studies. OR: Odds Ratio. CI: Confidence interval.

**Supplementary Table 3**
**Genetic association of rs13128814 with cardiovascular traits/diseases.** The summary statistics of genetic association of rs13128814 with 9 cardiovascular traits/diseases is indicated. SCAD: Spontaneous coronary artery dissection. CAD: Coronary artery disease. FMD: Fibromuscular dysplasia. SBP: Systolic blood pressure. DBP: Diastolic blood pressure. PP: Pulse pressure. AAdia: Diameter of ascending aorta. AAmax: Maximum area of ascending aorta. AAmin: Minimum area of ascending aorta.

**Supplementary Table 4**
**Prediction for rs13128814-dependent transcription factor binding sites.** Differential affinity of rs13128814 alleles with transcription factor binding sites from HOCOMOCO database (v12) were estimated using PERFECTOS-APE package. Approximate P-values for both variants were estimated using threshold method [48].

**Supplementary Table 5**
**Differential gene expression after *ZNF827* knockdown.** Mean expression, Log2 Fold Change, P-value and adjusted P-value were estimated using DESeq2 package, taking into account all samples, samples from BJ fibroblasts and samples from iPSC-derived SMCs. All genes with adjusted P-value below 0.05 taking into account all samples are indicated.

**Supplementary Table 6**
**Gene ontology enrichment in *ZNF827* target genes.** Enrichment of gene ontology terms for 429 consistent ZNF827 target genes was estimated using clusterprofiler R package.

**Supplementary Table 1**

| rsID | Coord (hg19) | Alleles | MAF | Distance | Dprime | R2 | Correlated Alleles | FORGEdb | Regulome | Function |
|---|---|---|---|---|---|---|---|---|---|---|
| rs1507928 | chr4:146788035 | (T/C) | 0.45 | 0 | 1 | 1 | T=T,C=C | 7 | 5 | NA |
| rs11435300 | chr4:146789510 | (-/T) | 0.44 | 1475 | 1 | 0.9761 | T=-,C=T | NA | . | NA |
| rs33934805 | chr4:146796087 | (A/G) | 0.44 | 8052 | 1 | 0.9761 | T=A,C=G | 7 | 7 | NA |
| rs28590383 | chr4:146803248 | (T/C) | 0.44 | 15213 | 0.9959 | 0.9761 | T=T,C=C | 10 | 4 | NA |
| rs10006310 | chr4:146809998 | (T/G) | 0.44 | 21963 | 0.9959 | 0.9761 | T=T,C=G | 6 | 4 | NA |
| rs10024759 | chr4:146797538 | (A/G) | 0.44 | 9503 | 1 | 0.9722 | T=A,C=G | 7 | 5 | NA |
| rs13124853 | chr4:146784774 | (A/G) | 0.44 | -3261 | 1 | 0.9683 | T=A,C=G | 7 | 3a | NA |
| rs1979974 | chr4:146800815 | (A/G) | 0.44 | 12780 | 1 | 0.9683 | T=A,C=G | 8 | 4 | NA |
| rs17020769 | chr4:146800922 | (C/T) | 0.44 | 12887 | 1 | 0.9683 | T=C,C=T | 9 | 4 | NA |
| rs10003835 | chr4:146809991 | (A/G) | 0.45 | 21956 | 0.9798 | 0.9524 | T=A,C=G | 6 | 4 | NA |
| rs7662070 | chr4:146809017 | (T/C) | 0.46 | 20982 | 0.9793 | 0.8958 | T=T,C=C | 9 | 4 | NA |
| rs7662069 | chr4:146809016 | (T/G) | 0.47 | 20981 | 0.9792 | 0.8921 | T=T,C=G | 9 | 4 | NA |
| rs4835266 | chr4:146821725 | (T/C) | 0.46 | 33690 | 0.9628 | 0.8729 | T=T,C=C | 4 | 6 | NA |
| rs10000888 | chr4:146809168 | (A/G) | 0.49 | 21133 | 0.9775 | 0.7521 | T=A,C=G | 9 | 4 | NA |
| rs7666150 | chr4:146814640 | (T/C) | 0.50 | 26605 | 0.9731 | 0.7513 | T=T,C=C | 6 | 5 | NA |
| rs7679068 | chr4:146808682 | (C/T) | 0.49 | 20647 | 0.973 | 0.7482 | T=C,C=T | 8 | 4 | NA |
| rs11100902 | chr4:146795226 | (A/G) | 0.48 | 7191 | 0.977 | 0.7191 | T=A,C=G | 7 | 5 | NA |
| rs13128814 | chr4:146801002 | (G/A) | 0.48 | 12967 | 0.9678 | 0.7085 | T=G,C=A | 10 | 2b | NA |
| rs11376412 | chr4:146776737 | (-/C) | 0.48 | -11298 | 0.9487 | 0.662 | T=-,C=C | NA | . | NA |
| rs988163 | chr4:146764883 | (C/T) | 0.48 | -23152 | 0.9311 | 0.6585 | T=C,C=T | 9 | 3a | NA |
| rs12646870 | chr4:146764120 | (T/C) | 0.48 | -23915 | 0.9311 | 0.6585 | T=T,C=C | 9 | 6 | NA |
| rs4835259 | chr4:146760042 | (T/G) | 0.48 | -27993 | 0.9311 | 0.6585 | T=T,C=G | 6 | 6 | NA |
| rs10004823 | chr4:146759607 | (C/T) | 0.48 | -28428 | 0.9311 | 0.6585 | T=C,C=T | 7 | 5 | NA |
| rs13120596 | chr4:146775541 | (G/A) | 0.47 | -12494 | 0.9387 | 0.6354 | T=G,C=A | 7 | 5 | NA |
| rs4345206 | chr4:146766536 | (T/C) | 0.47 | -21499 | 0.9296 | 0.6281 | T=T,C=C | 8 | 5 | NA |
| rs4835022 | chr4:146756490 | (T/C) | 0.47 | -31545 | 0.925 | 0.6244 | T=T,C=C | NA | 7 | NA |
| rs56080563 | chr4:146764376 | (-/A) | 0.47 | -23659 | 0.7693 | 0.5398 | T=-,C=A | NA | . | NA |
| rs6847076 | chr4:146758040 | (G/A) | 0.47 | -29995 | 0.7465 | 0.5185 | T=G,C=A | 6 | 6 | NA |

**Supplementary Table 2**

| Disease/Trait | Chr | Pos (hg38) | Risk allele | Lead SNP | P | OR or BETA | 95% CI description | PMID |
|---|---|---|---|---|---|---|---|---|
| Height | 4 | 145879770 | T | rs17020769 | 2.0E-75 | 0.0113 | [0.01-0.012] unit increase | 36224396 |
| Systolic blood pressure | 4 | 145900573 | ? | rs4835266 | 8.0E-17 | NA | | 30595370 |
| Pulse pressure | 4 | 145879770 | T | rs17020769 | 2.0E-11 | 0.0122 | [0.0087-0.0157] unit decrease | 34594039 |
| Alanine aminotransferase levels | 4 | 145879663 | G | rs1979974 | 7.0E-10 | 0.0218141 | [0.015-0.029] unit increase | 34594039 |
| Ascending aorta minimum area | 4 | 145879770 | C | rs17020769 | 7.0E-10 | 0.0448731 | [0.031-0.058] decrease | 34968759 |
| LDL cholesterol levels x short total sleep time interaction (2df test) | 4 | 145887530 | ? | rs7679068 | 2.0E-09 | 0.4651 | [0.13-0.8] unit increase | 31719535 |
| Coronary artery disease | 4 | 145888846 | ? | rs10006310 | 3.0E-09 | 0.0290661 | [0.019-0.039] unit decrease | 33020668 |
| Alcohol-associated liver disease in light drinkers | 4 | 145876386 | ? | rs10024759 | 3.0E-09 | 1.273 | [1.175-1.38] | 34387878 |
| Ascending aorta maximum area | 4 | 145879663 | A | rs1979974 | 4.0E-09 | 0.0451114 | [0.03-0.06] decrease | 34968759 |
| Ascending aorta diameter | 4 | 145874074 | G | rs11100902 | 6.0E-09 | 0.04 | unit increase | 35637384 |
| Spontaneous coronary artery dissection | 4 | 145866883 | C | rs1507928 | 9.0E-09 | 1.25 | [1.16-1.35] | 37248441 |
| Ascending aorta distensibility (MTAG) | 4 | 145879770 | C | rs17020769 | 4.0E-08 | 0.03 | unit increase | 35922433 |
| Ascending aorta maximum area (MTAG) | 4 | 145879850 | G | rs13128814 | 4.0E-08 | 6.56 | unit decrease | 35922433 |
| Systolic blood pressure (standard GWA) | 4 | 145879850 | G | rs13128814 | 4.0E-08 | 0.2882 | [0.19-0.39] unit increase | 37106081 |

**Supplementary Table 3**

| rsID | CHR | POS (hg19) | REF | ALT | Disease | BETA | SE | P |
|---|---|---|---|---|---|---|---|---|
| rs13128814 | 4 | 146801002 | G | A | SCAD | 0.216 | 0.039 | 3.2E-08 |
| | | | | | CAD | -0.027 | 0.005 | 5.9E-08 |
| | | | | | FMD | 0.148 | 0.044 | 7.7E-04 |
| | | | | | PP | -0.156 | 0.021 | 4.2E-14 |
| | | | | | DBP | -0.064 | 0.018 | 2.5E-04 |
| | | | | | SBP | -0.217 | 0.031 | 1.2E-12 |
| | | | | | AAdia | 0.042 | 0.007 | 7.0E-09 |
| | | | | | AAmax | 0.045 | 0.007 | 6.9E-09 |
| | | | | | AAmin | 0.045 | 0.007 | 6.8E-10 |

**Supplementary Table 4**

| Motif | Gene name | allele 1/allele 2 | P-value 1 | P-value 2 | Fold change | Direction |
|---|---|---|---|---|---|---|
| NFIA.H12INVIVO.0.P.B | NFIA | G/A | 5.5E-03 | 6.82E-05 | 81 | UP |
| NFIC.H12INVIVO.1.PSM.A | NFIC | G/A | 5.8E-03 | 8.20E-05 | 71 | UP |
| NR6A1.H12INVIVO.0.P.C | NR6A1 | G/A | 3.7E-03 | 8.34E-05 | 45 | UP |
| NFIC.H12INVIVO.0.PSM.A | NFIC | G/A | 3.7E-03 | 1.23E-04 | 30 | UP |
| NR6A1.H12INVIVO.1.P.C | NR6A1 | G/A | 9.9E-03 | 4.17E-04 | 24 | UP |
| NFIX.H12INVIVO.1.S.D | NFIX | G/A | 5.7E-03 | 2.67E-04 | 21 | UP |
| NFIB.H12INVIVO.1.PS.A | NFIB | G/A | 4.6E-03 | 2.31E-04 | 20 | UP |
| ZN554.H12INVIVO.1.P.C | ZNF554 | G/A | 5.59E-04 | 3.43E-05 | 16 | UP |
| ZBT43.H12INVIVO.0.S.D | ZBTB43 | G/A | 3.7E-03 | 4.59E-04 | 8 | UP |
| ZN649.H12INVIVO.0.P.C | ZNF649 | G/A | 9.19E-05 | 1.18E-05 | 8 | UP |
| SPI1.H12INVIVO.0.P.B | SPI1 | G/A | 1.30E-05 | 7.94E-05 | 6 | DOWN |
| ZN205.H12INVIVO.0.P.C | ZNF205 | G/A | 8.26E-04 | 1.43E-04 | 6 | UP |

**Supplementary Table 5**

| | | | All samples | | | BJ fibroblasts | | | iPSC-derived SMCs | | |
|---|---|---|---|---|---|---|---|---|---|---|---|
| **Ensembl Gene ID** | **Gene name** | **Mean expression** | **Log2 Fold Change** | **P** | **P.adj** | **Log2 Fold Change** | **P** | **P.adj** | **Log2 Fold Change** | **P** | **P.adj** |
| ENSG00000205250 | *E2F4* | 1268 | 0.9 | 2.1E-28 | 1.6E-24 | 1.2 | 9.9E-37 | 6.8E-34 | 0.7 | 4.3E-33 | 1.2E-29 |
| ENSG00000106443 | *PHF14* | 370 | 1.3 | 1.3E-28 | 1.6E-24 | 1.4 | 9.8E-20 | 1.3E-17 | 1.3 | 4.4E-28 | 7.4E-25 |
| ENSG00000175203 | *DCTN2* | 3983 | 0.4 | 2.5E-24 | 1.3E-20 | 0.5 | 2.6E-10 | 1.0E-08 | 0.4 | 3.5E-12 | 8.6E-10 |
| ENSG00000172340 | *SUCLG2* | 948 | 0.9 | 5.2E-24 | 2.0E-20 | 0.9 | 1.1E-21 | 1.9E-19 | 0.8 | 6.7E-30 | 1.4E-26 |
| ENSG00000163683 | *SMIM14* | 746 | 1.0 | 3.9E-21 | 1.2E-17 | 1.0 | 7.9E-22 | 1.4E-19 | 1.0 | 1.9E-20 | 1.0E-17 |
| ENSG00000145354 | *CISD2* | 399 | 1.0 | 5.7E-20 | 1.5E-16 | 1.2 | 2.6E-21 | 4.2E-19 | 0.9 | 2.1E-14 | 6.6E-12 |
| ENSG00000188554 | *NBR1* | 2234 | 1.0 | 8.3E-20 | 1.8E-16 | 1.5 | 1.6E-75 | 5.6E-72 | 0.7 | 3.5E-24 | 3.6E-21 |
| ENSG00000185127 | *C6orf120* | 973 | 0.9 | 9.8E-20 | 1.8E-16 | 1.2 | 8.8E-28 | 3.2E-25 | 0.7 | 3.9E-19 | 1.9E-16 |
| ENSG00000133059 | *DSTYK* | 866 | 0.7 | 1.0E-19 | 1.8E-16 | 0.9 | 1.7E-18 | 2.0E-16 | 0.5 | 1.0E-12 | 2.8E-10 |
| ENSG00000054983 | *GALC* | 368 | 0.8 | 1.3E-19 | 2.0E-16 | 0.8 | 6.6E-08 | 1.6E-06 | 0.7 | 1.9E-16 | 8.2E-14 |
| ENSG00000136026 | *CKAP4* | 12239 | 1.7 | 5.9E-19 | 8.2E-16 | 2.7 | 2.1E-123 | 2.9E-119 | 1.4 | 1.6E-73 | 2.1E-69 |
| ENSG00000099256 | *PRTFDC1* | 351 | 0.8 | 4.1E-18 | 5.3E-15 | 1.1 | 1.6E-16 | 1.5E-14 | 0.7 | 6.8E-11 | 1.4E-08 |
| ENSG00000150764 | *DIXDC1* | 751 | 1.1 | 4.7E-18 | 5.5E-15 | 1.6 | 5.8E-26 | 1.6E-23 | 0.9 | 5.5E-24 | 5.4E-21 |
| ENSG00000105197 | *TIMM50* | 1160 | 0.9 | 1.5E-17 | 1.7E-14 | 1.3 | 6.1E-43 | 5.3E-40 | 0.8 | 3.9E-19 | 1.9E-16 |
| ENSG00000001629 | *ANKIB1* | 1063 | 0.8 | 6.8E-17 | 7.0E-14 | 1.2 | 3.9E-24 | 8.7E-22 | 0.7 | 6.6E-21 | 4.1E-18 |
| ENSG00000048392 | *RRM2B* | 996 | 0.9 | 1.7E-16 | 1.7E-13 | 1.4 | 1.9E-27 | 6.5E-25 | 0.7 | 5.9E-23 | 4.9E-20 |
| ENSG00000182534 | *MXRA7* | 5968 | 0.7 | 3.5E-16 | 3.1E-13 | 0.9 | 1.8E-27 | 6.4E-25 | 0.6 | 8.7E-35 | 2.9E-31 |
| ENSG00000100889 | *PCK2* | 1484 | 0.6 | 3.6E-16 | 3.1E-13 | 0.5 | 1.2E-08 | 3.5E-07 | 0.6 | 4.3E-13 | 1.2E-10 |
| ENSG00000167323 | *STIM1* | 1371 | 0.8 | 8.5E-16 | 6.8E-13 | 1.1 | 3.5E-27 | 1.1E-24 | 0.7 | 7.1E-30 | 1.4E-26 |
| ENSG00000102096 | *PIM2* | 233 | 0.7 | 1.5E-15 | 1.1E-12 | 0.6 | 9.2E-05 | 9.9E-04 | 0.7 | 1.9E-13 | 5.7E-11 |
| ENSG00000141232 | *TOB1* | 1046 | 0.6 | 4.1E-15 | 3.0E-12 | 1.0 | 2.0E-21 | 3.3E-19 | 0.5 | 3.5E-11 | 8.6E-09 |
| ENSG00000188725 | *SMIM15* | 705 | 0.8 | 1.5E-14 | 1.0E-11 | 1.1 | 5.7E-14 | 4.1E-12 | 0.7 | 3.5E-13 | 1.0E-10 |
| ENSG00000106609 | *TMEM248* | 2453 | 0.5 | 1.7E-14 | 1.1E-11 | 0.9 | 2.2E-24 | 5.1E-22 | 0.4 | 8.9E-16 | 3.5E-13 |
| ENSG00000132824 | *SERINC3* | 2824 | 0.8 | 2.0E-14 | 1.3E-11 | 1.0 | 2.7E-28 | 1.0E-25 | 0.7 | 3.3E-24 | 3.6E-21 |
| ENSG00000182004 | *SNRPE* | 701 | 0.9 | 8.2E-14 | 5.0E-11 | 1.3 | 7.9E-21 | 1.2E-18 | 0.7 | 5.2E-15 | 1.9E-12 |
| ENSG00000188021 | *UBQLN2* | 1543 | 0.8 | 1.3E-13 | 7.9E-11 | 1.3 | 2.5E-34 | 1.5E-31 | 0.6 | 1.9E-13 | 5.7E-11 |
| ENSG00000151892 | *GFRA1* | 3346 | 0.6 | 1.5E-13 | 8.5E-11 | 0.3 | 3.0E-04 | 2.8E-03 | 0.7 | 2.9E-21 | 2.0E-18 |
| ENSG00000174099 | *MSRB3* | 4706 | -0.4 | 2.4E-13 | 1.3E-10 | -0.2 | 1.7E-03 | 1.1E-02 | -0.5 | 7.9E-21 | 4.6E-18 |
| ENSG00000109756 | *RAPGEF2* | 524 | 0.8 | 1.2E-12 | 6.4E-10 | 0.7 | 3.1E-09 | 1.1E-07 | 0.9 | 2.2E-15 | 8.4E-13 |
| ENSG00000100567 | *PSMA3* | 1274 | 0.8 | 2.0E-12 | 1.0E-09 | 0.9 | 6.7E-17 | 6.4E-15 | 0.8 | 3.0E-21 | 2.0E-18 |
| ENSG00000138386 | *NAB1* | 386 | 0.8 | 3.8E-12 | 1.9E-09 | 1.1 | 4.1E-13 | 2.5E-11 | 0.6 | 2.3E-09 | 3.7E-07 |
| ENSG00000134874 | *DZIP1* | 741 | 0.6 | 4.2E-12 | 2.0E-09 | 0.8 | 5.4E-15 | 4.4E-13 | 0.5 | 5.0E-11 | 1.1E-08 |
| ENSG00000057663 | *ATG5* | 541 | 0.8 | 5.1E-12 | 2.4E-09 | 1.3 | 3.5E-30 | 1.6E-27 | 0.6 | 7.0E-10 | 1.3E-07 |
| ENSG00000101577 | *LPIN2* | 925 | 0.9 | 8.0E-12 | 3.6E-09 | 1.6 | 2.0E-53 | 4.0E-50 | 0.8 | 8.1E-22 | 6.1E-19 |
| ENSG00000126107 | *HECTD3* | 2009 | -0.4 | 2.3E-11 | 9.9E-09 | -0.4 | 6.1E-07 | 1.2E-05 | -0.3 | 1.2E-07 | 1.4E-05 |
| ENSG00000151612 | *ZNF827* | 272 | 1.9 | 6.4E-11 | 2.7E-08 | 2.2 | 3.1E-59 | 7.2E-56 | 1.9 | 4.2E-10 | 7.7E-08 |
| ENSG00000274211 | *SOCS7* | 467 | -0.5 | 7.0E-11 | 2.9E-08 | -0.6 | 1.0E-06 | 1.9E-05 | -0.4 | 1.7E-06 | 1.3E-04 |
| ENSG00000101782 | *RIOK3* | 1337 | -0.4 | 7.6E-11 | 3.0E-08 | -0.5 | 1.8E-07 | 4.1E-06 | -0.3 | 5.4E-07 | 5.2E-05 |
| ENSG00000154124 | *OTULIN* | 517 | 0.5 | 9.9E-11 | 3.9E-08 | 0.5 | 2.7E-05 | 3.4E-04 | 0.4 | 2.4E-07 | 2.5E-05 |
| ENSG00000105974 | *CAV1* | 20568 | 0.5 | 3.2E-10 | 1.2E-07 | 0.6 | 1.0E-14 | 7.6E-13 | 0.5 | 3.2E-17 | 1.4E-14 |
| ENSG00000169583 | *CLIC3* | 282 | -0.8 | 3.3E-10 | 1.2E-07 | -0.7 | 8.2E-10 | 3.1E-08 | -0.9 | 6.2E-11 | 1.3E-08 |
| ENSG00000153179 | *RASSF3* | 2635 | -0.4 | 3.7E-10 | 1.4E-07 | -0.5 | 1.6E-07 | 3.6E-06 | -0.3 | 2.0E-09 | 3.5E-07 |
| ENSG00000160208 | *RRP1B* | 905 | 0.5 | 4.7E-10 | 1.7E-07 | 0.7 | 2.2E-10 | 9.2E-09 | 0.4 | 1.6E-06 | 1.2E-04 |
| ENSG00000125834 | *STK35* | 830 | -0.5 | 6.2E-10 | 2.2E-07 | -0.5 | 3.5E-09 | 1.2E-07 | -0.4 | 8.4E-09 | 1.2E-06 |
| ENSG00000170445 | *HARS1* | 1758 | 0.7 | 8.0E-10 | 2.7E-07 | 0.9 | 1.1E-23 | 2.2E-21 | 0.6 | 1.2E-24 | 1.7E-21 |
| ENSG00000041357 | *PSMA4* | 2112 | 0.5 | 9.1E-10 | 3.0E-07 | 0.7 | 6.1E-14 | 4.3E-12 | 0.4 | 2.7E-09 | 4.2E-07 |
| ENSG00000115935 | *WIPF1* | 1050 | 0.7 | 1.0E-09 | 3.3E-07 | 1.1 | 1.0E-32 | 5.1E-30 | 0.5 | 6.8E-08 | 7.7E-06 |
| ENSG00000181467 | *RAP2B* | 1239 | 0.6 | 1.1E-09 | 3.4E-07 | 1.2 | 1.1E-33 | 6.0E-31 | 0.4 | 1.3E-10 | 2.5E-08 |
| ENSG00000114999 | *TTL* | 2237 | -0.3 | 1.4E-09 | 4.4E-07 | -0.2 | 3.7E-02 | 1.3E-01 | -0.4 | 3.7E-11 | 8.7E-09 |
| ENSG00000137992 | *DBT* | 402 | 0.5 | 1.7E-09 | 5.2E-07 | 0.3 | 4.4E-03 | 2.5E-02 | 0.6 | 6.9E-13 | 1.9E-10 |
| ENSG00000198700 | *IPO9* | 2504 | 0.5 | 2.9E-09 | 8.5E-07 | 1.0 | 2.0E-29 | 8.2E-27 | 0.3 | 2.4E-09 | 3.8E-07 |
| ENSG00000081087 | *OSTM1* | 519 | 0.6 | 3.0E-09 | 8.9E-07 | 0.9 | 6.5E-13 | 3.9E-11 | 0.4 | 2.6E-06 | 1.9E-04 |
| ENSG00000148943 | *LIN7C* | 388 | 0.6 | 3.1E-09 | 9.0E-07 | 0.9 | 4.0E-09 | 1.3E-07 | 0.4 | 2.2E-06 | 1.6E-04 |
| ENSG00000238227 | *TMEM250* | 1751 | -0.4 | 3.7E-09 | 1.0E-06 | -0.4 | 7.2E-08 | 1.8E-06 | -0.3 | 5.7E-07 | 5.3E-05 |
| ENSG00000173011 | *TADA2B* | 939 | -0.5 | 4.2E-09 | 1.2E-06 | -0.6 | 1.7E-08 | 4.8E-07 | -0.3 | 1.6E-06 | 1.2E-04 |
| ENSG00000124535 | *WRNIP1* | 778 | 0.2 | 4.2E-09 | 1.2E-06 | 0.2 | 1.3E-02 | 5.9E-02 | 0.2 | 5.1E-05 | 2.2E-03 |
| ENSG00000047634 | *SCML1* | 224 | 0.6 | 1.1E-08 | 2.8E-06 | 0.7 | 4.6E-05 | 5.5E-04 | 0.5 | 1.5E-05 | 8.5E-04 |
| ENSG00000158985 | *CDC42SE2* | 755 | 0.8 | 1.2E-08 | 3.2E-06 | 0.9 | 1.4E-10 | 6.2E-09 | 0.9 | 1.1E-20 | 6.0E-18 |
| ENSG00000153310 | *CYRIB* | 763 | -0.5 | 1.3E-08 | 3.3E-06 | -0.6 | 6.1E-10 | 2.3E-08 | -0.4 | 4.4E-09 | 6.6E-07 |
| ENSG00000180921 | *FAM83H* | 968 | -0.7 | 1.5E-08 | 3.8E-06 | -1.3 | 4.9E-24 | 1.1E-21 | -0.5 | 5.6E-07 | 5.3E-05 |
| ENSG00000205339 | *IPO7* | 4359 | 0.6 | 1.5E-08 | 3.8E-06 | 0.6 | 7.4E-09 | 2.3E-07 | 0.5 | 2.1E-10 | 3.9E-08 |
| ENSG00000152700 | *SAR1B* | 1300 | 0.5 | 2.2E-08 | 5.5E-06 | 0.8 | 4.7E-19 | 5.9E-17 | 0.3 | 5.6E-05 | 2.4E-03 |
| ENSG00000075785 | *RAB7A* | 5918 | 0.6 | 2.5E-08 | 6.0E-06 | 1.1 | 1.2E-47 | 1.7E-44 | 0.3 | 4.3E-11 | 9.9E-09 |
| ENSG00000143390 | *RFX5* | 751 | 0.4 | 2.5E-08 | 6.0E-06 | 0.5 | 7.5E-08 | 1.8E-06 | 0.2 | 1.1E-04 | 4.0E-03 |
| ENSG00000138071 | *ACTR2* | 7756 | 0.6 | 2.7E-08 | 6.2E-06 | 1.0 | 7.3E-28 | 2.7E-25 | 0.5 | 2.2E-13 | 6.5E-11 |
| ENSG00000144136 | *SLC20A1* | 1532 | 0.7 | 3.9E-08 | 9.0E-06 | 1.5 | 2.9E-42 | 2.3E-39 | 0.5 | 6.9E-15 | 2.4E-12 |
| ENSG00000152952 | *PLOD2* | 3057 | 0.4 | 4.3E-08 | 9.8E-06 | 0.7 | 5.0E-12 | 2.6E-10 | 0.3 | 1.3E-05 | 7.5E-04 |

| | | | | | | | | | | | |
|---|---|---|---|---|---|---|---|---|---|---|---|
| ENSG00000106070 | *GRB10* | 1326 | -0.4 | 4.6E-08 | 1.0E-05 | -0.5 | 1.1E-09 | 4.1E-08 | -0.2 | 1.7E-04 | 5.6E-03 |
| ENSG00000198252 | *STYX* | 554 | 0.7 | 4.9E-08 | 1.1E-05 | 0.9 | 4.5E-09 | 1.5E-07 | 0.6 | 7.1E-08 | 7.9E-06 |
| ENSG00000160714 | *UBE2Q1* | 1947 | 0.3 | 5.3E-08 | 1.2E-05 | 0.5 | 2.4E-08 | 6.6E-07 | 0.3 | 7.1E-06 | 4.4E-04 |
| ENSG00000119487 | *MAPKAP1* | 3124 | -0.3 | 5.7E-08 | 1.2E-05 | -0.2 | 2.4E-03 | 1.5E-02 | -0.3 | 1.1E-06 | 9.2E-05 |
| ENSG00000198915 | *RASGEF1A* | 38 | -1.1 | 5.7E-08 | 1.2E-05 | -0.7 | 9.4E-04 | 6.9E-03 | -0.7 | 1.8E-05 | 9.7E-04 |
| ENSG00000164761 | *TNFRSF11B* | 3458 | 0.9 | 6.0E-08 | 1.2E-05 | 1.8 | 2.8E-95 | 1.9E-91 | 0.5 | 5.0E-06 | 3.4E-04 |
| ENSG00000139921 | *TMX1* | 663 | 0.9 | 6.0E-08 | 1.2E-05 | 2.2 | 3.3E-79 | 1.5E-75 | 0.6 | 3.6E-15 | 1.4E-12 |
| ENSG00000070610 | *GBA2* | 1098 | -0.4 | 6.4E-08 | 1.3E-05 | -0.9 | 4.1E-15 | 3.5E-13 | -0.3 | 1.7E-05 | 9.3E-04 |
| ENSG00000166619 | *BLCAP* | 1962 | -0.4 | 6.9E-08 | 1.4E-05 | -0.6 | 2.3E-14 | 1.7E-12 | -0.3 | 9.8E-06 | 5.8E-04 |
| ENSG00000152193 | *OBI1* | 262 | 0.8 | 6.9E-08 | 1.4E-05 | 0.9 | 2.2E-07 | 4.8E-06 | 0.8 | 7.2E-11 | 1.5E-08 |
| ENSG00000006607 | *FARP2* | 1327 | -0.3 | 7.0E-08 | 1.4E-05 | -0.3 | 7.6E-04 | 5.9E-03 | -0.2 | 2.5E-05 | 1.2E-03 |
| ENSG00000143337 | *TOR1AIP1* | 1304 | 0.3 | 7.1E-08 | 1.4E-05 | 0.4 | 1.7E-07 | 3.8E-06 | 0.3 | 9.0E-07 | 7.9E-05 |
| ENSG00000276293 | *PIP4K2B* | 3622 | -0.2 | 8.8E-08 | 1.7E-05 | -0.2 | 2.7E-03 | 1.7E-02 | -0.3 | 3.8E-09 | 5.9E-07 |
| ENSG00000173210 | *ABLIM3* | 803 | -0.5 | 9.4E-08 | 1.7E-05 | -0.4 | 2.4E-07 | 5.1E-06 | -0.3 | 7.7E-04 | 1.8E-02 |
| ENSG00000111196 | *MAGOHB* | 293 | 0.6 | 9.4E-08 | 1.7E-05 | 0.7 | 2.5E-06 | 4.2E-05 | 0.6 | 5.3E-07 | 5.1E-05 |
| ENSG00000143622 | *RIT1* | 1073 | -0.3 | 9.5E-08 | 1.7E-05 | -0.4 | 3.1E-06 | 5.2E-05 | -0.2 | 1.8E-04 | 5.8E-03 |
| ENSG00000153885 | *KCTD15* | 1284 | 0.2 | 9.2E-08 | 1.7E-05 | 0.3 | 7.7E-04 | 5.9E-03 | 0.2 | 2.2E-04 | 7.0E-03 |
| ENSG00000013588 | *GPRC5A* | 1950 | -0.5 | 1.1E-07 | 1.9E-05 | -0.7 | 4.6E-10 | 1.8E-08 | -0.4 | 1.4E-08 | 1.9E-06 |
| ENSG00000198730 | *CTR9* | 1050 | 0.5 | 1.2E-07 | 2.2E-05 | 0.7 | 6.4E-10 | 2.4E-08 | 0.4 | 1.3E-07 | 1.4E-05 |
| ENSG00000180398 | *MCFD2* | 2992 | 0.6 | 1.3E-07 | 2.2E-05 | 1.1 | 5.9E-45 | 6.3E-42 | 0.3 | 1.7E-08 | 2.1E-06 |
| ENSG00000165097 | *KDM1B* | 369 | 0.8 | 1.3E-07 | 2.3E-05 | 1.6 | 8.4E-31 | 4.0E-28 | 0.5 | 1.6E-08 | 2.1E-06 |
| ENSG00000111707 | *SUDS3* | 888 | 0.5 | 1.4E-07 | 2.5E-05 | 0.7 | 2.6E-12 | 1.4E-10 | 0.4 | 2.1E-07 | 2.2E-05 |
| ENSG00000120705 | *ETF1* | 3267 | 0.2 | 1.6E-07 | 2.6E-05 | 0.2 | 6.7E-03 | 3.4E-02 | 0.3 | 1.8E-07 | 1.9E-05 |
| ENSG00000131876 | *SNRPA1* | 806 | -0.3 | 1.6E-07 | 2.6E-05 | -0.3 | 2.4E-03 | 1.5E-02 | -0.3 | 6.6E-05 | 2.7E-03 |
| ENSG00000144369 | *FAM171B* | 186 | 0.7 | 2.1E-07 | 3.5E-05 | 0.7 | 3.9E-08 | 1.0E-06 | 0.4 | 2.4E-04 | 7.5E-03 |
| ENSG00000140632 | *GLYR1* | 2529 | -0.2 | 2.5E-07 | 4.0E-05 | -0.3 | 1.7E-04 | 1.7E-03 | -0.2 | 9.9E-05 | 3.7E-03 |
| ENSG00000168256 | *NKIRAS2* | 2661 | -0.2 | 2.5E-07 | 4.1E-05 | -0.4 | 3.1E-08 | 8.4E-07 | -0.1 | 5.4E-04 | 1.4E-02 |
| ENSG00000271601 | *LIX1L* | 1787 | -0.2 | 2.9E-07 | 4.6E-05 | -0.2 | 2.0E-03 | 1.3E-02 | -0.2 | 4.4E-05 | 2.0E-03 |
| ENSG00000151725 | *CENPU* | 297 | 0.6 | 3.4E-07 | 5.4E-05 | 1.2 | 1.7E-09 | 6.1E-08 | 0.4 | 3.1E-05 | 1.5E-03 |
| ENSG00000134265 | *NAPG* | 657 | 0.3 | 3.5E-07 | 5.5E-05 | 0.3 | 3.3E-04 | 3.0E-03 | 0.3 | 1.1E-04 | 4.0E-03 |
| ENSG00000104435 | *STMN2* | 921 | -0.6 | 3.7E-07 | 5.7E-05 | -1.1 | 6.2E-33 | 3.3E-30 | -0.3 | 3.0E-04 | 8.9E-03 |
| ENSG00000141376 | *BCAS3* | 447 | 0.5 | 3.7E-07 | 5.7E-05 | 0.8 | 1.8E-09 | 6.3E-08 | 0.4 | 1.8E-08 | 2.2E-06 |
| ENSG00000170653 | *ATF7* | 850 | 0.4 | 5.1E-07 | 7.7E-05 | 0.5 | 1.3E-07 | 3.0E-06 | 0.3 | 1.8E-04 | 5.8E-03 |
| ENSG00000006576 | *PHTF2* | 1674 | 0.5 | 5.4E-07 | 8.1E-05 | 0.8 | 1.7E-11 | 8.0E-10 | 0.4 | 2.3E-09 | 3.7E-07 |
| ENSG00000102349 | *KLF8* | 71 | -0.7 | 5.6E-07 | 8.3E-05 | -0.4 | 4.4E-03 | 2.5E-02 | -0.5 | 6.9E-05 | 2.8E-03 |
| ENSG00000074696 | *HACD3* | 3253 | 0.5 | 6.0E-07 | 9.0E-05 | 0.8 | 3.9E-16 | 3.5E-14 | 0.4 | 9.9E-10 | 1.7E-07 |
| ENSG00000197903 | *H2BC12* | 1061 | -0.5 | 6.3E-07 | 9.2E-05 | -1.0 | 5.3E-22 | 9.2E-20 | -0.3 | 1.6E-06 | 1.2E-04 |
| ENSG00000171488 | *LRRC8C* | 201 | 0.7 | 6.5E-07 | 9.2E-05 | 1.3 | 2.6E-18 | 3.0E-16 | 0.4 | 6.2E-05 | 2.6E-03 |
| ENSG00000184787 | *UBE2G2* | 1798 | 0.3 | 6.4E-07 | 9.2E-05 | 0.6 | 1.2E-11 | 5.8E-10 | 0.2 | 3.6E-05 | 1.7E-03 |
| ENSG00000109670 | *FBXW7* | 253 | 0.4 | 6.4E-07 | 9.2E-05 | 0.2 | 1.0E-01 | 2.6E-01 | 0.5 | 1.1E-06 | 9.4E-05 |
| ENSG00000088930 | *XRN2* | 2052 | 0.5 | 7.1E-07 | 1.0E-04 | 1.0 | 1.0E-24 | 2.5E-22 | 0.3 | 5.4E-09 | 8.0E-07 |
| ENSG00000171174 | *RBKS* | 82 | -0.7 | 8.0E-07 | 1.1E-04 | -1.3 | 5.2E-13 | 3.1E-11 | -0.2 | 9.4E-03 | 1.1E-01 |
| ENSG00000124523 | *SIRT5* | 203 | 0.5 | 8.2E-07 | 1.1E-04 | 0.7 | 1.8E-06 | 3.1E-05 | 0.2 | 1.3E-03 | 2.7E-02 |
| ENSG00000142494 | *SLC47A1* | 128 | 0.6 | 8.9E-07 | 1.2E-04 | 0.3 | 2.4E-02 | 9.0E-02 | 0.5 | 5.2E-06 | 3.5E-04 |
| ENSG00000169446 | *MMGT1* | 736 | 0.6 | 9.4E-07 | 1.3E-04 | 1.2 | 3.3E-18 | 3.8E-16 | 0.4 | 4.7E-06 | 3.2E-04 |
| ENSG00000134375 | *TIMM17A* | 1449 | 0.4 | 9.4E-07 | 1.3E-04 | 0.8 | 1.6E-16 | 1.5E-14 | 0.3 | 1.8E-05 | 9.7E-04 |
| ENSG00000122884 | *P4HA1* | 2413 | 0.2 | 1.0E-06 | 1.3E-04 | 0.2 | 2.9E-03 | 1.7E-02 | 0.2 | 3.5E-05 | 1.7E-03 |
| ENSG00000116285 | *ERRFI1* | 3987 | 0.5 | 1.1E-06 | 1.5E-04 | 0.8 | 1.2E-10 | 5.2E-09 | 0.4 | 1.6E-07 | 1.8E-05 |
| ENSG00000061987 | *MON2* | 697 | 0.3 | 1.1E-06 | 1.5E-04 | 0.5 | 3.1E-07 | 6.4E-06 | 0.2 | 1.2E-03 | 2.5E-02 |
| ENSG00000135341 | *MAP3K7* | 1143 | 0.3 | 1.2E-06 | 1.6E-04 | 0.5 | 2.7E-09 | 9.3E-08 | 0.2 | 1.7E-03 | 3.2E-02 |
| ENSG00000170027 | *YWHAG* | 5295 | 0.6 | 1.3E-06 | 1.6E-04 | 1.3 | 3.6E-53 | 6.3E-50 | 0.3 | 2.1E-09 | 3.6E-07 |
| ENSG00000164938 | *TP53INP1* | 861 | 0.6 | 1.4E-06 | 1.8E-04 | 0.8 | 5.7E-08 | 1.4E-06 | 0.7 | 1.7E-12 | 4.4E-10 |
| ENSG00000012174 | *MBTPS2* | 666 | 0.5 | 1.4E-06 | 1.8E-04 | 0.9 | 1.3E-19 | 1.7E-17 | 0.2 | 3.8E-04 | 1.1E-02 |
| ENSG00000005189 | *REXO5* | 125 | 0.6 | 1.4E-06 | 1.8E-04 | 1.1 | 6.7E-08 | 1.7E-06 | 0.2 | 2.5E-03 | 4.1E-02 |
| ENSG00000118276 | *B4GALT6* | 522 | 0.9 | 1.5E-06 | 1.9E-04 | 0.4 | 7.4E-03 | NA | 1.0 | 1.9E-22 | 1.5E-19 |
| ENSG00000165092 | *ALDH1A1* | 11155 | 0.3 | 1.6E-06 | 2.0E-04 | 0.2 | 8.9E-04 | 6.6E-03 | 0.3 | 5.7E-11 | 1.2E-08 |
| ENSG00000161981 | *SNRNP25* | 1077 | 0.3 | 1.6E-06 | 2.0E-04 | 0.4 | 6.3E-06 | 9.6E-05 | 0.2 | 6.0E-04 | 1.5E-02 |
| ENSG00000030110 | *BAK1* | 1316 | -0.2 | 2.0E-06 | 2.5E-04 | -0.3 | 2.2E-03 | 1.4E-02 | -0.2 | 2.8E-04 | 8.3E-03 |
| ENSG00000078177 | *N4BP2* | 179 | 0.7 | 2.1E-06 | 2.5E-04 | 0.8 | 2.3E-05 | 3.0E-04 | 0.5 | 2.0E-05 | 1.0E-03 |
| ENSG00000143549 | *TPM3* | 11754 | -0.2 | 2.2E-06 | 2.6E-04 | -0.1 | 4.0E-02 | 1.3E-01 | -0.2 | 9.7E-07 | 8.4E-05 |
| ENSG00000164104 | *HMGB2* | 1393 | 0.5 | 2.3E-06 | 2.7E-04 | 1.1 | 2.9E-27 | 9.5E-25 | 0.2 | 1.1E-05 | 6.4E-04 |
| ENSG00000117461 | *PIK3R3* | 766 | 0.5 | 2.5E-06 | 3.0E-04 | 0.6 | 5.0E-05 | 5.9E-04 | 0.4 | 1.1E-12 | 2.9E-10 |
| ENSG00000107819 | *SFXN3* | 3100 | -0.3 | 2.5E-06 | 3.0E-04 | -0.5 | 1.0E-09 | 3.8E-08 | -0.2 | 2.4E-04 | 7.3E-03 |
| ENSG00000205189 | *ZBTB10* | 323 | 0.5 | 2.5E-06 | 3.0E-04 | 0.5 | 1.7E-04 | 1.7E-03 | 0.4 | 1.9E-05 | 1.0E-03 |
| ENSG00000138434 | *ITPRID2* | 2591 | -0.3 | 2.9E-06 | 3.4E-04 | -0.5 | 9.2E-08 | 2.2E-06 | -0.2 | 1.1E-06 | 9.2E-05 |
| ENSG00000126391 | *FRMD8* | 1159 | -0.2 | 3.1E-06 | 3.5E-04 | -0.1 | 6.0E-02 | 1.8E-01 | -0.2 | 1.0E-04 | 3.8E-03 |
| ENSG00000139926 | *FRMD6* | 7566 | -0.2 | 3.1E-06 | 3.5E-04 | -0.1 | 1.3E-01 | 3.1E-01 | -0.3 | 2.6E-09 | 4.2E-07 |
| ENSG00000178607 | *ERN1* | 121 | 0.5 | 3.9E-06 | 4.5E-04 | 0.3 | 1.4E-02 | 6.1E-02 | 0.3 | 4.3E-04 | 1.2E-02 |
| ENSG00000112305 | *SMAP1* | 850 | -0.3 | 4.0E-06 | 4.5E-04 | -0.2 | 4.5E-03 | 2.5E-02 | -0.3 | 7.1E-05 | 2.8E-03 |
| ENSG00000156639 | *ZFAND3* | 3448 | -0.2 | 4.5E-06 | 5.1E-04 | -0.2 | 9.6E-04 | 7.1E-03 | -0.2 | 4.9E-05 | 2.2E-03 |
| ENSG00000139636 | *LMBR1L* | 843 | -0.5 | 5.3E-06 | 5.9E-04 | -1.2 | 7.0E-27 | 2.2E-24 | -0.2 | 6.5E-04 | 1.6E-02 |
| ENSG00000113712 | *CSNK1A1* | 3711 | -0.2 | 6.2E-06 | 6.9E-04 | -0.3 | 1.9E-04 | 1.8E-03 | -0.1 | 6.3E-03 | 8.1E-02 |
| ENSG00000125651 | *GTF2F1* | 2344 | 0.3 | 6.3E-06 | 6.9E-04 | 0.4 | 2.7E-05 | 3.5E-04 | 0.3 | 5.7E-05 | 2.4E-03 |
| ENSG00000197594 | *ENPP1* | 793 | 0.3 | 6.7E-06 | 7.3E-04 | 0.5 | 1.5E-07 | 3.3E-06 | 0.2 | 1.2E-03 | 2.6E-02 |

| | | | | | | | | | | | |
|---|---|---|---|---|---|---|---|---|---|---|---|
| ENSG00000112902 | *SEMA5A* | 8193 | 0.2 | 6.9E-06 | 7.4E-04 | 0.2 | 2.8E-03 | 1.7E-02 | 0.3 | 1.4E-05 | 8.0E-04 |
| ENSG00000166579 | *NDEL1* | 1550 | -0.2 | 7.1E-06 | 7.5E-04 | -0.3 | 4.2E-04 | 3.6E-03 | -0.1 | 1.5E-02 | 1.4E-01 |
| ENSG00000197296 | *FITM2* | 394 | 0.6 | 7.3E-06 | 7.7E-04 | 1.2 | 1.9E-22 | 3.3E-20 | 0.2 | 1.9E-03 | 3.4E-02 |
| ENSG00000196182 | *STK40* | 3633 | -0.5 | 7.3E-06 | 7.7E-04 | -1.2 | 2.8E-44 | 2.8E-41 | -0.2 | 2.0E-05 | 1.1E-03 |
| ENSG00000134686 | *PHC2* | 2795 | -0.1 | 7.5E-06 | 7.9E-04 | -0.2 | 2.9E-02 | 1.1E-01 | -0.1 | 2.3E-03 | 3.9E-02 |
| ENSG00000076067 | *RBMS2* | 2492 | -0.3 | 7.8E-06 | 8.1E-04 | -0.4 | 3.2E-06 | 5.2E-05 | -0.2 | 1.5E-05 | 8.5E-04 |
| ENSG00000146386 | *ABRACL* | 645 | -0.4 | 8.0E-06 | 8.3E-04 | -0.7 | 1.6E-12 | 9.0E-11 | -0.2 | 2.4E-03 | 4.1E-02 |
| ENSG00000174839 | *DENND6A* | 256 | 0.4 | 8.6E-06 | 8.9E-04 | 0.4 | 2.1E-03 | 1.3E-02 | 0.3 | 6.1E-05 | 2.6E-03 |
| ENSG00000117069 | *ST6GALNAC[illegible]* | 502 | 0.4 | 8.7E-06 | 8.9E-04 | 0.4 | 1.2E-02 | 5.3E-02 | 0.3 | 2.9E-06 | 2.0E-04 |
| ENSG00000116396 | *KCNC4* | 509 | -0.5 | 8.9E-06 | 9.0E-04 | -0.3 | 2.5E-03 | 1.5E-02 | -0.5 | 5.1E-06 | 3.4E-04 |
| ENSG00000169760 | *NLGN1* | 255 | 0.3 | 9.1E-06 | 9.2E-04 | 0.4 | 2.1E-04 | 2.0E-03 | 0.2 | 2.4E-03 | 4.1E-02 |
| ENSG00000104472 | *CHRAC1* | 747 | 0.3 | 9.5E-06 | 9.4E-04 | 0.6 | 2.5E-07 | 5.3E-06 | 0.2 | 3.0E-03 | 4.8E-02 |
| ENSG00000183722 | *LHFPL6* | 1276 | 0.5 | 1.0E-05 | 9.9E-04 | 1.3 | 3.7E-43 | 3.5E-40 | 0.2 | 6.1E-05 | 2.5E-03 |
| ENSG00000173852 | *DPY19L1* | 910 | 0.6 | 1.2E-05 | 1.2E-03 | 1.3 | 1.6E-23 | 3.3E-21 | 0.3 | 5.4E-06 | 3.5E-04 |
| ENSG00000103381 | *CPPED1* | 1013 | -0.3 | 1.2E-05 | 1.2E-03 | -0.2 | 2.3E-02 | 8.8E-02 | -0.4 | 1.0E-06 | 8.6E-05 |
| ENSG00000171241 | *SHCBP1* | 957 | 0.7 | 1.2E-05 | 1.2E-03 | 1.9 | 5.1E-49 | 7.9E-46 | 0.4 | 6.6E-09 | 9.5E-07 |
| ENSG00000055211 | *GINM1* | 1004 | 0.4 | 1.4E-05 | 1.3E-03 | 0.7 | 4.3E-12 | 2.3E-10 | 0.2 | 5.5E-04 | 1.4E-02 |
| ENSG00000197622 | *CDC42SE1* | 1720 | -0.2 | 1.5E-05 | 1.4E-03 | -0.5 | 5.6E-09 | 1.8E-07 | -0.1 | 2.5E-03 | 4.2E-02 |
| ENSG00000100228 | *RAB36* | 328 | -0.2 | 1.5E-05 | 1.4E-03 | -0.2 | 2.7E-02 | 9.8E-02 | -0.2 | 2.1E-03 | 3.7E-02 |
| ENSG00000267106 | *ZNF561-AS1* | 153 | 0.4 | 1.6E-05 | 1.5E-03 | 0.2 | 2.7E-02 | 1.0E-01 | 0.4 | 1.7E-04 | 5.7E-03 |
| ENSG00000149357 | *LAMTOR1* | 3400 | 0.3 | 1.6E-05 | 1.5E-03 | 0.7 | 1.2E-14 | 8.6E-13 | 0.2 | 1.7E-04 | 5.6E-03 |
| ENSG00000077713 | *SLC25A43* | 608 | -0.3 | 1.8E-05 | 1.7E-03 | -0.2 | 1.4E-02 | 6.0E-02 | -0.3 | 1.8E-04 | 5.8E-03 |
| ENSG00000085788 | *DDHD2* | 761 | 0.2 | 1.8E-05 | 1.7E-03 | 0.2 | 2.6E-02 | 9.8E-02 | 0.2 | 2.6E-04 | 7.7E-03 |
| ENSG00000100441 | *KHNYN* | 2100 | -0.4 | 1.9E-05 | 1.8E-03 | -0.9 | 4.3E-26 | 1.2E-23 | -0.2 | 9.2E-05 | 3.5E-03 |
| ENSG00000139645 | *ANKRD52* | 6485 | -0.4 | 2.1E-05 | 1.9E-03 | -0.9 | 7.2E-30 | 3.1E-27 | -0.2 | 3.7E-03 | 5.6E-02 |
| ENSG00000132768 | *DPH2* | 894 | -0.2 | 2.2E-05 | 2.0E-03 | -0.3 | 3.3E-03 | 1.9E-02 | -0.2 | 1.2E-03 | 2.4E-02 |
| ENSG00000137273 | *FOXF2* | 867 | 0.2 | 2.3E-05 | 2.1E-03 | 0.1 | 1.5E-01 | 3.3E-01 | 0.3 | 2.3E-06 | 1.7E-04 |
| ENSG00000063978 | *RNF4* | 1999 | -0.2 | 2.4E-05 | 2.2E-03 | -0.2 | 3.8E-03 | 2.2E-02 | -0.1 | 6.0E-03 | 7.7E-02 |
| ENSG00000165169 | *DYNLT3* | 607 | -0.4 | 2.5E-05 | 2.2E-03 | -0.9 | 4.8E-10 | 1.9E-08 | -0.3 | 6.7E-06 | 4.2E-04 |
| ENSG00000114251 | *WNT5A* | 5737 | 0.3 | 2.5E-05 | 2.2E-03 | 0.8 | 1.6E-19 | 2.1E-17 | 0.2 | 5.6E-03 | 7.4E-02 |
| ENSG00000154319 | *FAM167A* | 308 | 0.6 | 2.6E-05 | 2.3E-03 | 1.0 | 2.7E-24 | 6.1E-22 | 0.1 | 5.2E-02 | NA |
| ENSG00000128607 | *KLHDC10* | 1406 | 0.5 | 2.7E-05 | 2.4E-03 | 0.5 | 1.1E-06 | 2.0E-05 | 0.6 | 2.4E-23 | 2.1E-20 |
| ENSG00000135720 | *DYNC1LI2* | 2988 | -0.2 | 2.9E-05 | 2.5E-03 | -0.4 | 3.0E-07 | 6.2E-06 | -0.2 | 4.0E-04 | 1.1E-02 |
| ENSG00000236609 | *ZNF853* | 170 | 0.5 | 2.9E-05 | 2.5E-03 | 0.6 | 6.8E-04 | 5.4E-03 | 0.4 | 2.1E-04 | 6.6E-03 |
| ENSG00000162946 | *DISC1* | 78 | 0.5 | 3.0E-05 | 2.6E-03 | 0.9 | 2.9E-05 | 3.6E-04 | 0.2 | 8.3E-03 | 9.8E-02 |
| ENSG00000196459 | *TRAPPC2* | 364 | -0.3 | 3.0E-05 | 2.6E-03 | -0.1 | 1.4E-01 | 3.2E-01 | -0.3 | 2.3E-05 | 1.2E-03 |
| ENSG00000197747 | *S100A10* | 11206 | -0.3 | 3.1E-05 | 2.7E-03 | -0.7 | 1.1E-20 | 1.6E-18 | -0.2 | 4.5E-05 | 2.0E-03 |
| ENSG00000110906 | *KCTD10* | 4599 | -0.1 | 3.2E-05 | 2.7E-03 | -0.2 | 6.0E-03 | 3.1E-02 | -0.1 | 3.5E-03 | 5.3E-02 |
| ENSG00000170473 | *PYM1* | 651 | -0.3 | 3.2E-05 | 2.7E-03 | -0.6 | 2.8E-08 | 7.7E-07 | -0.2 | 5.5E-03 | 7.3E-02 |
| ENSG00000164649 | *CDCA7L* | 821 | 0.2 | 3.2E-05 | 2.7E-03 | 0.3 | 4.3E-04 | 3.7E-03 | 0.2 | 8.5E-04 | 1.9E-02 |
| ENSG00000119638 | *NEK9* | 2186 | 0.2 | 3.2E-05 | 2.7E-03 | 0.2 | 3.0E-02 | 1.1E-01 | 0.1 | 1.9E-03 | 3.5E-02 |
| ENSG00000145022 | *TCTA* | 1927 | -0.2 | 3.4E-05 | 2.8E-03 | -0.2 | 2.5E-03 | 1.5E-02 | -0.1 | 4.3E-03 | 6.1E-02 |
| ENSG00000149485 | *FADS1* | 5821 | 0.2 | 3.4E-05 | 2.9E-03 | 0.5 | 2.5E-08 | 6.8E-07 | 0.1 | 3.5E-03 | 5.3E-02 |
| ENSG00000180198 | *RCC1* | 1050 | 0.2 | 3.5E-05 | 2.9E-03 | 0.3 | 5.3E-04 | 4.4E-03 | 0.1 | 4.3E-03 | 6.1E-02 |
| ENSG00000136273 | *HUS1* | 497 | -0.2 | 3.5E-05 | 2.9E-03 | -0.4 | 5.0E-05 | 5.8E-04 | -0.1 | 1.8E-02 | 1.6E-01 |
| ENSG00000131873 | *CHSY1* | 1224 | 0.3 | 3.6E-05 | 3.0E-03 | 0.8 | 3.1E-14 | 2.2E-12 | 0.1 | 3.3E-03 | 5.2E-02 |
| ENSG00000172059 | *KLF11* | 577 | -0.2 | 3.7E-05 | 3.0E-03 | -0.3 | 1.5E-03 | 1.0E-02 | -0.2 | 1.1E-03 | 2.4E-02 |
| ENSG00000060642 | *PIGV* | 318 | 0.2 | 3.7E-05 | 3.0E-03 | 0.3 | 4.9E-03 | 2.7E-02 | 0.2 | 2.1E-03 | 3.7E-02 |
| ENSG00000181827 | *RFX7* | 482 | 0.3 | 3.8E-05 | 3.1E-03 | 0.2 | 6.1E-02 | 1.8E-01 | 0.3 | 6.1E-05 | 2.5E-03 |
| ENSG00000165802 | *NSMF* | 1937 | -0.2 | 3.9E-05 | 3.1E-03 | -0.1 | 8.4E-02 | 2.2E-01 | -0.3 | 6.6E-06 | 4.2E-04 |
| ENSG00000166822 | *TMEM170A* | 130 | 0.6 | 3.9E-05 | 3.1E-03 | 0.6 | 2.6E-04 | 2.4E-03 | 0.5 | 1.0E-04 | 3.7E-03 |
| ENSG00000009335 | *UBE3C* | 4012 | -0.2 | 4.0E-05 | 3.2E-03 | -0.3 | 6.1E-04 | 4.9E-03 | -0.1 | 5.2E-03 | 7.0E-02 |
| ENSG00000198324 | *PHETA1* | 555 | -0.2 | 4.1E-05 | 3.2E-03 | -0.2 | 7.6E-03 | 3.8E-02 | -0.2 | 3.5E-03 | 5.4E-02 |
| ENSG00000131263 | *RLIM* | 1050 | 0.3 | 4.2E-05 | 3.3E-03 | 0.5 | 1.3E-06 | 2.4E-05 | 0.2 | 4.2E-03 | 6.1E-02 |
| ENSG00000134287 | *ARF3* | 5327 | -0.2 | 4.3E-05 | 3.3E-03 | -0.3 | 1.9E-04 | 1.8E-03 | -0.1 | 1.4E-04 | 4.7E-03 |
| ENSG00000121039 | *RDH10* | 3182 | 0.6 | 4.4E-05 | 3.4E-03 | 1.6 | 9.9E-69 | 2.7E-65 | 0.7 | 6.9E-47 | 4.7E-43 |
| ENSG00000183044 | *ABAT* | 249 | 0.6 | 4.6E-05 | 3.6E-03 | 1.2 | 1.0E-10 | 4.5E-09 | 0.6 | 1.2E-08 | 1.6E-06 |
| ENSG00000164040 | *PGRMC2* | 1438 | 0.2 | 4.6E-05 | 3.6E-03 | 0.2 | 1.1E-02 | 4.9E-02 | 0.2 | 1.8E-04 | 5.7E-03 |
| ENSG00000232320 | *MXRA7P1* | 14 | 0.6 | 4.7E-05 | 3.6E-03 | 0.2 | 1.2E-01 | NA | 0.2 | 1.2E-03 | NA |
| ENSG00000137478 | *FCHSD2* | 933 | 0.2 | 4.7E-05 | 3.6E-03 | 0.6 | 6.0E-10 | 2.3E-08 | 0.1 | 2.3E-02 | 1.8E-01 |
| ENSG00000146729 | *NIPSNAP2* | 1007 | 0.3 | 4.8E-05 | 3.6E-03 | 0.5 | 1.2E-06 | 2.2E-05 | 0.2 | 3.0E-03 | 4.9E-02 |
| ENSG00000100731 | *PCNX1* | 1001 | 0.3 | 4.9E-05 | 3.7E-03 | 0.4 | 4.9E-05 | 5.8E-04 | 0.2 | 2.1E-03 | 3.7E-02 |
| ENSG00000166471 | *TMEM41B* | 318 | 0.3 | 5.1E-05 | 3.8E-03 | 0.7 | 2.9E-08 | 7.9E-07 | 0.1 | 1.7E-02 | 1.5E-01 |
| ENSG00000109046 | *WSB1* | 740 | -0.4 | 5.2E-05 | 3.9E-03 | -0.3 | 2.8E-03 | 1.7E-02 | -0.6 | 6.7E-15 | 2.4E-12 |
| ENSG00000108578 | *BLMH* | 1542 | -0.2 | 5.4E-05 | 4.0E-03 | -0.2 | 4.3E-03 | 2.4E-02 | -0.1 | 5.7E-03 | 7.5E-02 |
| ENSG00000093217 | *XYLB* | 85 | 0.5 | 5.5E-05 | 4.1E-03 | 0.9 | 4.8E-06 | 7.6E-05 | 0.1 | 1.7E-02 | 1.5E-01 |
| ENSG00000155304 | *HSPA13* | 852 | 0.3 | 5.6E-05 | 4.1E-03 | 0.6 | 1.2E-09 | 4.4E-08 | 0.1 | 1.1E-02 | 1.2E-01 |
| ENSG00000265972 | *TXNIP* | 3024 | -0.4 | 6.0E-05 | 4.4E-03 | -0.9 | 3.7E-25 | 9.4E-23 | -0.2 | 6.7E-04 | 1.6E-02 |
| ENSG00000125870 | *SNRPB2* | 1126 | 0.3 | 6.4E-05 | 4.7E-03 | 0.3 | 1.4E-04 | 1.5E-03 | 0.4 | 5.6E-06 | 3.6E-04 |
| ENSG00000143493 | *INTS7* | 633 | 0.2 | 6.7E-05 | 4.8E-03 | 0.2 | 6.4E-03 | 3.3E-02 | 0.1 | 7.3E-03 | 8.9E-02 |
| ENSG00000079974 | *RABL2B* | 398 | 0.3 | 6.9E-05 | 5.0E-03 | 0.2 | 2.1E-02 | 8.1E-02 | 0.3 | 1.5E-04 | 5.1E-03 |
| ENSG00000143815 | *LBR* | 1342 | 0.4 | 7.1E-05 | 5.1E-03 | 1.1 | 5.1E-20 | 7.0E-18 | 0.2 | 1.9E-04 | 6.1E-03 |
| ENSG00000136710 | *CCDC115* | 641 | 0.2 | 7.1E-05 | 5.1E-03 | 0.6 | 2.0E-07 | 4.4E-06 | 0.1 | 1.4E-02 | 1.3E-01 |
| ENSG00000150403 | *TMCO3* | 2581 | 0.2 | 7.4E-05 | 5.3E-03 | 0.2 | 3.7E-03 | 2.1E-02 | 0.1 | 1.3E-02 | 1.3E-01 |

| | | | | | | | | | | | |
|---|---|---|---|---|---|---|---|---|---|---|---|
| ENSG00000038210 | *PI4K2B* | 680 | 0.5 | 7.7E-05 | 5.4E-03 | 1.1 | 3.0E-16 | 2.7E-14 | 0.5 | 5.7E-07 | 5.3E-05 |
| ENSG00000154889 | *MPPE1* | 364 | 0.3 | 7.7E-05 | 5.4E-03 | 0.4 | 3.0E-05 | 3.7E-04 | 0.3 | 6.5E-05 | 2.6E-03 |
| ENSG00000089902 | *RCOR1* | 1479 | -0.2 | 7.6E-05 | 5.4E-03 | -0.2 | 3.0E-03 | 1.8E-02 | -0.2 | 2.2E-04 | 6.7E-03 |
| ENSG00000135472 | *FAIM2* | 277 | 0.5 | 7.8E-05 | 5.4E-03 | 1.2 | 7.2E-08 | 1.8E-06 | 0.3 | 1.8E-05 | 9.9E-04 |
| ENSG00000100426 | *ZBED4* | 966 | -0.2 | 8.1E-05 | 5.6E-03 | -0.4 | 8.1E-06 | 1.2E-04 | -0.1 | 6.7E-03 | 8.3E-02 |
| ENSG00000155111 | *CDK19* | 555 | -0.2 | 8.2E-05 | 5.6E-03 | -0.4 | 3.8E-05 | 4.6E-04 | -0.1 | 2.2E-02 | 1.7E-01 |
| ENSG00000184445 | *KNTC1* | 273 | 0.4 | 8.3E-05 | 5.7E-03 | 0.9 | 4.7E-09 | 1.5E-07 | 0.1 | 1.3E-02 | 1.3E-01 |
| ENSG00000103994 | *ZNF106* | 1584 | 0.3 | 8.6E-05 | 5.9E-03 | 0.1 | 1.6E-01 | 3.5E-01 | 0.4 | 1.3E-06 | 1.1E-04 |
| ENSG00000142875 | *PRKACB* | 1558 | -0.2 | 8.6E-05 | 5.9E-03 | -0.3 | 1.8E-03 | 1.2E-02 | -0.2 | 2.9E-04 | 8.5E-03 |
| ENSG00000117758 | *STX12* | 1902 | -0.2 | 9.2E-05 | 6.2E-03 | -0.2 | 3.3E-02 | 1.2E-01 | -0.2 | 7.2E-05 | 2.8E-03 |
| ENSG00000108179 | *PPIF* | 2561 | -0.2 | 9.3E-05 | 6.3E-03 | -0.6 | 5.8E-12 | 3.0E-10 | -0.1 | 1.2E-02 | 1.2E-01 |
| ENSG00000166847 | *DCTN5* | 3065 | -0.2 | 9.9E-05 | 6.7E-03 | -0.5 | 2.2E-10 | 9.1E-09 | -0.1 | 2.4E-03 | 4.0E-02 |
| ENSG00000164442 | *CITED2* | 15888 | 0.1 | 9.9E-05 | 6.7E-03 | 0.1 | 4.3E-02 | 1.4E-01 | 0.1 | 1.4E-02 | 1.3E-01 |
| ENSG00000181104 | *F2R* | 3519 | 0.2 | 1.0E-04 | 6.9E-03 | 0.5 | 4.1E-07 | 8.3E-06 | 0.1 | 8.0E-03 | 9.5E-02 |
| ENSG00000134153 | *EMC7* | 1310 | 0.2 | 1.0E-04 | 6.9E-03 | 0.4 | 3.3E-06 | 5.4E-05 | 0.2 | 7.3E-04 | 1.7E-02 |
| ENSG00000110756 | *HPS5* | 777 | 0.3 | 1.1E-04 | 7.1E-03 | 0.6 | 1.2E-08 | 3.6E-07 | 0.2 | 5.3E-04 | 1.4E-02 |
| ENSG00000137817 | *PARP6* | 494 | -0.2 | 1.1E-04 | 7.3E-03 | -0.3 | 8.1E-04 | 6.2E-03 | -0.1 | 1.6E-02 | 1.5E-01 |
| ENSG00000183853 | *KIRREL1* | 5298 | 0.3 | 1.1E-04 | 7.4E-03 | 0.6 | 4.7E-08 | 1.2E-06 | 0.2 | 1.2E-03 | 2.4E-02 |
| ENSG00000128973 | *CLN6* | 366 | 0.3 | 1.1E-04 | 7.4E-03 | 0.9 | 3.0E-09 | 1.0E-07 | 0.1 | 1.3E-02 | 1.3E-01 |
| ENSG00000116406 | *EDEM3* | 621 | 0.3 | 1.1E-04 | 7.5E-03 | 0.6 | 1.4E-08 | 4.1E-07 | 0.2 | 3.6E-03 | 5.4E-02 |
| ENSG00000146676 | *PURB* | 1466 | 0.2 | 1.2E-04 | 7.5E-03 | 0.2 | 6.5E-02 | 1.9E-01 | 0.4 | 1.1E-08 | 1.4E-06 |
| ENSG00000198830 | *HMGN2* | 6539 | 0.2 | 1.2E-04 | 7.7E-03 | 0.5 | 1.8E-11 | 8.5E-10 | 0.1 | 5.5E-03 | 7.3E-02 |
| ENSG00000135929 | *CYP27A1* | 325 | 0.3 | 1.2E-04 | 7.7E-03 | 0.6 | 3.2E-07 | 6.5E-06 | 0.1 | 2.9E-02 | 2.1E-01 |
| ENSG00000111371 | *SLC38A1* | 3209 | 0.3 | 1.3E-04 | 8.4E-03 | 0.3 | 4.6E-04 | 3.9E-03 | 0.4 | 8.1E-11 | 1.6E-08 |
| ENSG00000122140 | *MRPS2* | 1482 | -0.2 | 1.4E-04 | 8.6E-03 | -0.2 | 1.4E-02 | 6.0E-02 | -0.2 | 9.6E-05 | 3.6E-03 |
| ENSG00000185344 | *ATP6V0A2* | 545 | 0.3 | 1.4E-04 | 9.0E-03 | 0.6 | 1.7E-05 | 2.3E-04 | 0.2 | 3.9E-04 | 1.1E-02 |
| ENSG00000183520 | *UTP11* | 700 | 0.2 | 1.4E-04 | 9.0E-03 | 0.2 | 3.6E-02 | 1.2E-01 | 0.2 | 1.8E-03 | 3.3E-02 |
| ENSG00000085978 | *ATG16L1* | 659 | -0.2 | 1.5E-04 | 9.3E-03 | -0.4 | 2.9E-05 | 3.6E-04 | -0.1 | 3.9E-02 | 2.4E-01 |
| ENSG00000103005 | *USB1* | 2063 | 0.3 | 1.5E-04 | 9.5E-03 | 0.9 | 1.0E-22 | 1.8E-20 | 0.1 | 1.3E-02 | 1.3E-01 |
| ENSG00000153904 | *DDAH1* | 7726 | -0.3 | 1.5E-04 | 9.6E-03 | -0.9 | 1.1E-13 | 7.2E-12 | -0.2 | 1.1E-04 | 4.0E-03 |
| ENSG00000102908 | *NFAT5* | 849 | 0.4 | 1.5E-04 | 9.6E-03 | 0.4 | 3.3E-05 | 4.1E-04 | 0.5 | 1.0E-06 | 8.8E-05 |
| ENSG00000145332 | *KLHL8* | 337 | 0.3 | 1.7E-04 | 1.0E-02 | 0.7 | 4.1E-07 | 8.3E-06 | 0.1 | 1.8E-02 | 1.6E-01 |
| ENSG00000147601 | *TERF1* | 796 | 0.4 | 1.7E-04 | 1.0E-02 | 0.9 | 1.2E-12 | 7.1E-11 | 0.5 | 1.1E-09 | 1.8E-07 |
| ENSG00000175376 | *EIF1AD* | 729 | -0.2 | 1.7E-04 | 1.0E-02 | -0.1 | 1.5E-01 | 3.3E-01 | -0.2 | 1.3E-03 | 2.7E-02 |
| ENSG00000185361 | *TNFAIP8L1* | 228 | 0.3 | 1.8E-04 | 1.1E-02 | 0.8 | 6.1E-07 | 1.2E-05 | 0.1 | 2.9E-02 | 2.1E-01 |
| ENSG00000182199 | *SHMT2* | 6053 | -0.1 | 1.8E-04 | 1.1E-02 | -0.2 | 4.0E-04 | 3.5E-03 | -0.1 | 2.2E-02 | 1.8E-01 |
| ENSG00000198331 | *HYLS1* | 185 | 0.3 | 1.8E-04 | 1.1E-02 | 0.6 | 1.3E-05 | 1.9E-04 | 0.1 | 1.9E-02 | 1.6E-01 |
| ENSG00000147669 | *POLR2K* | 598 | 0.3 | 1.8E-04 | 1.1E-02 | 0.6 | 1.6E-08 | 4.6E-07 | 0.2 | 1.1E-02 | 1.1E-01 |
| ENSG00000107443 | *CCNJ* | 202 | 0.3 | 1.8E-04 | 1.1E-02 | 0.2 | 7.8E-02 | 2.1E-01 | 0.3 | 4.2E-05 | 1.9E-03 |
| ENSG00000136444 | *RSAD1* | 498 | -0.2 | 1.8E-04 | 1.1E-02 | -0.2 | 2.4E-02 | 9.2E-02 | -0.1 | 2.1E-02 | 1.7E-01 |
| ENSG00000147471 | *PLPBP* | 1262 | 0.2 | 1.8E-04 | 1.1E-02 | 0.3 | 2.0E-03 | 1.3E-02 | 0.2 | 1.1E-03 | 2.3E-02 |
| ENSG00000153786 | *ZDHHC7* | 2723 | -0.1 | 1.8E-04 | 1.1E-02 | -0.2 | 3.9E-03 | 2.2E-02 | -0.1 | 1.8E-02 | 1.5E-01 |
| ENSG00000140367 | *UBE2Q2* | 1591 | 0.2 | 1.9E-04 | 1.1E-02 | 0.3 | 1.5E-03 | 1.0E-02 | 0.2 | 2.1E-04 | 6.7E-03 |
| ENSG00000117519 | *CNN3* | 7967 | 0.2 | 1.9E-04 | 1.1E-02 | 0.3 | 1.4E-05 | 2.0E-04 | 0.1 | 7.9E-03 | 9.4E-02 |
| ENSG00000139436 | *GIT2* | 1399 | 0.2 | 1.9E-04 | 1.1E-02 | 0.3 | 5.5E-05 | 6.3E-04 | 0.2 | 4.0E-03 | 5.9E-02 |
| ENSG00000153130 | *SCOC* | 601 | 0.5 | 2.0E-04 | 1.2E-02 | 0.5 | 6.6E-03 | 3.4E-02 | 0.5 | 3.1E-08 | 3.8E-06 |
| ENSG00000141510 | *TP53* | 2499 | -0.1 | 2.0E-04 | 1.2E-02 | -0.2 | 2.2E-02 | 8.7E-02 | -0.1 | 5.8E-03 | 7.6E-02 |
| ENSG00000169762 | *TAPT1* | 241 | -0.3 | 2.0E-04 | 1.2E-02 | -0.1 | 2.0E-01 | 4.0E-01 | -0.5 | 9.1E-07 | 8.0E-05 |
| ENSG00000008283 | *CYB561* | 721 | -0.2 | 2.2E-04 | 1.3E-02 | -0.2 | 2.9E-03 | 1.7E-02 | -0.2 | 3.4E-04 | 9.5E-03 |
| ENSG00000129472 | *RAB2B* | 232 | 0.3 | 2.2E-04 | 1.3E-02 | 0.4 | 7.5E-04 | 5.8E-03 | 0.1 | 1.2E-02 | 1.2E-01 |
| ENSG00000159147 | *DONSON* | 340 | 0.2 | 2.3E-04 | 1.3E-02 | 0.5 | 6.2E-05 | 7.1E-04 | 0.1 | 4.0E-02 | 2.5E-01 |
| ENSG00000154920 | *EME1* | 66 | 0.4 | 2.3E-04 | 1.3E-02 | 0.8 | 3.3E-04 | 3.0E-03 | 0.1 | 2.8E-02 | 2.0E-01 |
| ENSG00000167105 | *TMEM92* | 66 | -0.4 | 2.4E-04 | 1.4E-02 | -0.3 | 4.1E-02 | 1.4E-01 | -0.2 | 4.0E-03 | 5.9E-02 |
| ENSG00000204054 | *LINC00963* | 473 | 0.2 | 2.4E-04 | 1.4E-02 | 0.2 | 1.4E-02 | 6.0E-02 | 0.3 | 7.4E-05 | 2.9E-03 |
| ENSG00000182636 | *NDN* | 1651 | 0.4 | 2.5E-04 | 1.4E-02 | 1.2 | 1.5E-26 | 4.6E-24 | 0.5 | 2.2E-19 | 1.1E-16 |
| ENSG00000137075 | *RNF38* | 989 | -0.3 | 2.7E-04 | 1.5E-02 | -1.1 | 2.5E-16 | 2.3E-14 | -0.2 | 1.4E-03 | 2.8E-02 |
| ENSG00000170088 | *TMEM192* | 501 | 0.3 | 2.7E-04 | 1.5E-02 | 0.7 | 2.5E-09 | 8.7E-08 | 0.3 | 1.2E-04 | 4.2E-03 |
| ENSG00000159259 | *CHAF1B* | 310 | 0.3 | 2.7E-04 | 1.5E-02 | 1.1 | 2.8E-10 | 1.1E-08 | 0.1 | 1.6E-02 | 1.4E-01 |
| ENSG00000119541 | *VPS4B* | 1110 | 0.2 | 2.8E-04 | 1.5E-02 | 0.4 | 1.3E-05 | 1.8E-04 | 0.2 | 7.0E-04 | 1.7E-02 |
| ENSG00000162694 | *EXTL2* | 474 | 0.3 | 2.9E-04 | 1.6E-02 | 0.9 | 8.7E-11 | 3.9E-09 | 0.1 | 2.1E-02 | 1.7E-01 |
| ENSG00000204604 | *ZNF468* | 199 | -0.2 | 2.9E-04 | 1.6E-02 | -0.2 | 5.0E-02 | 1.6E-01 | -0.1 | 1.4E-02 | 1.4E-01 |
| ENSG00000108669 | *CYTH1* | 301 | 0.3 | 2.9E-04 | 1.6E-02 | 0.2 | 6.2E-02 | 1.8E-01 | 0.4 | 1.7E-05 | 9.5E-04 |
| ENSG00000152684 | *PELO* | 1355 | 0.2 | 2.9E-04 | 1.6E-02 | 0.3 | 1.4E-03 | 9.5E-03 | 0.2 | 2.5E-04 | 7.7E-03 |
| ENSG00000172795 | *DCP2* | 709 | 0.4 | 3.0E-04 | 1.6E-02 | 0.4 | 5.0E-03 | 2.7E-02 | 1.0 | 4.9E-21 | 3.2E-18 |
| ENSG00000270170 | *NCBP2AS2* | 319 | -0.3 | 3.0E-04 | 1.6E-02 | -0.3 | 2.3E-03 | 1.5E-02 | -0.2 | 1.6E-03 | 3.1E-02 |
| ENSG00000136824 | *SMC2* | 355 | 0.3 | 3.0E-04 | 1.6E-02 | 0.8 | 2.4E-10 | 9.8E-09 | 0.2 | 3.1E-03 | 4.9E-02 |
| ENSG00000197951 | *ZNF71* | 354 | -0.2 | 3.0E-04 | 1.6E-02 | -0.4 | 2.1E-03 | 1.4E-02 | -0.1 | 2.5E-02 | 1.9E-01 |
| ENSG00000123395 | *ATG101* | 1503 | -0.2 | 3.1E-04 | 1.7E-02 | -0.3 | 8.9E-04 | 6.6E-03 | -0.1 | 1.1E-02 | 1.2E-01 |
| ENSG00000114626 | *ABTB1* | 477 | -0.2 | 3.2E-04 | 1.7E-02 | -0.3 | 2.8E-03 | 1.7E-02 | -0.2 | 4.2E-03 | 6.1E-02 |
| ENSG00000185963 | *BICD2* | 892 | 0.3 | 3.2E-04 | 1.7E-02 | 0.7 | 1.1E-09 | 3.9E-08 | 0.3 | 2.1E-05 | 1.1E-03 |
| ENSG00000272269 | *NUP153-AS1* | 64 | -0.4 | 3.3E-04 | 1.7E-02 | -0.2 | 1.6E-01 | 3.5E-01 | -0.3 | 8.5E-04 | 1.9E-02 |
| ENSG00000108639 | *SYNGR2* | 2426 | -0.2 | 3.3E-04 | 1.8E-02 | -0.3 | 8.5E-04 | 6.4E-03 | -0.3 | 4.6E-05 | 2.1E-03 |
| ENSG00000085840 | *ORC1* | 163 | 0.4 | 3.4E-04 | 1.8E-02 | 1.3 | 1.6E-08 | 4.7E-07 | 0.1 | 4.7E-02 | 2.7E-01 |
| ENSG00000147509 | *RGS20* | 68 | 0.3 | 3.4E-04 | 1.8E-02 | 0.2 | 1.3E-01 | 3.0E-01 | 0.2 | 8.6E-03 | 9.9E-02 |

| | | | | | | | | | | | |
|---|---|---|---|---|---|---|---|---|---|---|---|
| ENSG00000120802 | *TMPO* | 1722 | 0.3 | 3.5E-04 | 1.8E-02 | 1.0 | 1.3E-29 | 5.4E-27 | 0.1 | 2.3E-02 | 1.8E-01 |
| ENSG00000151468 | *CCDC3* | 5110 | -0.2 | 3.5E-04 | 1.8E-02 | 0.0 | 9.7E-01 | NA | -0.2 | 1.9E-05 | 1.0E-03 |
| ENSG00000163811 | *WDR43* | 965 | 0.2 | 3.5E-04 | 1.8E-02 | 0.2 | 3.3E-02 | 1.2E-01 | 0.2 | 1.7E-03 | 3.2E-02 |
| ENSG00000154856 | *APCDD1* | 1764 | 0.3 | 3.6E-04 | 1.8E-02 | 0.7 | 5.4E-16 | 4.8E-14 | 0.3 | 3.4E-08 | 4.1E-06 |
| ENSG00000227051 | *C14orf132* | 1076 | -0.2 | 3.6E-04 | 1.8E-02 | 0.0 | 7.2E-01 | 8.5E-01 | -0.5 | 1.5E-08 | 1.9E-06 |
| ENSG00000177426 | *TGIF1* | 645 | 0.2 | 3.6E-04 | 1.8E-02 | 0.2 | 1.4E-02 | 6.0E-02 | 0.1 | 8.6E-03 | 1.0E-01 |
| ENSG00000021355 | *SERPINB1* | 990 | 0.2 | 3.6E-04 | 1.9E-02 | 0.1 | 1.1E-01 | 2.8E-01 | 0.2 | 5.5E-04 | 1.4E-02 |
| ENSG00000256594 | *P11-705C15.* | 23 | -0.4 | 3.6E-04 | 1.9E-02 | -0.4 | 8.5E-03 | 4.1E-02 | -0.1 | 4.8E-02 | NA |
| ENSG00000036054 | *TBC1D23* | 911 | -0.2 | 3.8E-04 | 2.0E-02 | -0.4 | 4.5E-05 | 5.4E-04 | -0.2 | 5.8E-03 | 7.6E-02 |
| ENSG00000138594 | *TMOD3* | 2672 | -0.2 | 3.9E-04 | 2.0E-02 | -0.6 | 1.2E-10 | 5.2E-09 | -0.1 | 5.6E-03 | 7.4E-02 |
| ENSG00000077092 | *RARB* | 45 | 0.4 | 3.9E-04 | 2.0E-02 | 1.2 | 2.6E-08 | 7.1E-07 | 0.2 | 5.9E-03 | NA |
| ENSG00000104635 | *SLC39A14* | 2656 | 0.3 | 4.0E-04 | 2.0E-02 | 0.8 | 9.2E-19 | 1.1E-16 | 0.2 | 4.5E-05 | 2.0E-03 |
| ENSG00000211455 | *STK38L* | 1638 | -0.3 | 4.0E-04 | 2.0E-02 | -0.9 | 2.0E-12 | 1.1E-10 | -0.2 | 3.0E-04 | 8.8E-03 |
| ENSG00000174606 | *ANGEL2* | 421 | 0.3 | 4.0E-04 | 2.0E-02 | 0.7 | 9.8E-09 | 3.0E-07 | 0.3 | 2.6E-04 | 7.9E-03 |
| ENSG00000186166 | *CENATAC* | 188 | -0.3 | 4.0E-04 | 2.0E-02 | -0.9 | 4.0E-08 | 1.0E-06 | -0.1 | 2.1E-02 | 1.7E-01 |
| ENSG00000187837 | *H1-2* | 349 | -0.2 | 4.0E-04 | 2.0E-02 | -0.6 | 2.7E-07 | 5.7E-06 | -0.1 | 3.3E-02 | 2.2E-01 |
| ENSG00000215271 | *HOMEZ* | 453 | 0.2 | 4.1E-04 | 2.0E-02 | 0.1 | 3.5E-01 | 5.6E-01 | 0.2 | 1.9E-03 | 3.5E-02 |
| ENSG00000102034 | *ELF4* | 2116 | -0.2 | 4.2E-04 | 2.1E-02 | -0.5 | 1.1E-10 | 4.8E-09 | -0.1 | 5.7E-02 | 3.0E-01 |
| ENSG00000186594 | *MIR22HG* | 561 | -0.2 | 4.1E-04 | 2.1E-02 | -0.4 | 3.1E-06 | 5.2E-05 | -0.1 | 5.1E-02 | 2.8E-01 |
| ENSG00000116489 | *CAPZA1* | 4958 | -0.2 | 4.1E-04 | 2.1E-02 | -0.3 | 6.9E-05 | 7.7E-04 | -0.1 | 7.8E-03 | 9.3E-02 |
| ENSG00000275591 | *XKR5* | 43 | 0.4 | 4.2E-04 | 2.1E-02 | 1.7 | 3.5E-07 | 7.2E-06 | 0.1 | 8.5E-02 | 3.7E-01 |
| ENSG00000113389 | *NPR3* | 1205 | 0.2 | 4.2E-04 | 2.1E-02 | 0.3 | 1.7E-02 | 7.2E-02 | 0.3 | 1.6E-05 | 8.7E-04 |
| ENSG00000100578 | *KIAA0586* | 229 | 0.3 | 4.2E-04 | 2.1E-02 | 0.4 | 1.2E-03 | 8.3E-03 | 0.2 | 3.8E-03 | 5.7E-02 |
| ENSG00000072609 | *CHFR* | 1003 | -0.2 | 4.3E-04 | 2.1E-02 | -0.1 | 7.3E-02 | 2.0E-01 | -0.1 | 1.2E-03 | 2.4E-02 |
| ENSG00000099203 | *TMED1* | 625 | 0.2 | 4.4E-04 | 2.2E-02 | 0.3 | 2.3E-03 | 1.5E-02 | 0.1 | 7.1E-03 | 8.8E-02 |
| ENSG00000169718 | *DUS1L* | 1657 | -0.2 | 4.4E-04 | 2.2E-02 | -0.3 | 1.9E-04 | 1.9E-03 | -0.1 | 1.1E-02 | 1.2E-01 |
| ENSG00000132003 | *ZSWIM4* | 314 | -0.3 | 4.5E-04 | 2.2E-02 | -0.4 | 1.5E-05 | 2.1E-04 | -0.1 | 3.1E-02 | 2.1E-01 |
| ENSG00000100206 | *DMC1* | 23 | 0.4 | 4.6E-04 | 2.2E-02 | 1.4 | 4.3E-06 | 6.8E-05 | 0.1 | 9.8E-02 | NA |
| ENSG00000082146 | *STRADB* | 770 | 0.2 | 4.8E-04 | 2.3E-02 | 0.6 | 2.4E-06 | 4.1E-05 | 0.1 | 1.0E-02 | 1.1E-01 |
| ENSG00000121486 | *TRMT1L* | 443 | 0.2 | 4.8E-04 | 2.3E-02 | 0.4 | 7.2E-04 | 5.6E-03 | 0.1 | 8.1E-03 | 9.6E-02 |
| ENSG00000015475 | *BID* | 741 | -0.2 | 4.8E-04 | 2.3E-02 | -0.2 | 7.8E-03 | 3.9E-02 | -0.1 | 3.6E-02 | 2.3E-01 |
| ENSG00000164808 | *SPIDR* | 698 | 0.2 | 4.9E-04 | 2.3E-02 | 0.8 | 1.7E-15 | 1.4E-13 | 0.1 | 1.0E-01 | 4.0E-01 |
| ENSG00000104219 | *ZDHHC2* | 1039 | 0.2 | 4.9E-04 | 2.3E-02 | 0.3 | 7.4E-04 | 5.7E-03 | 0.2 | 1.7E-03 | 3.2E-02 |
| ENSG00000127022 | *CANX* | 16134 | 0.2 | 5.0E-04 | 2.4E-02 | 0.4 | 1.1E-08 | 3.5E-07 | 0.1 | 2.9E-02 | 2.1E-01 |
| ENSG00000211584 | *SLC48A1* | 915 | -0.2 | 5.0E-04 | 2.4E-02 | -0.5 | 3.1E-07 | 6.5E-06 | -0.1 | 4.2E-02 | 2.5E-01 |
| ENSG00000164647 | *STEAP1* | 39 | 0.4 | 5.0E-04 | 2.4E-02 | 0.8 | 2.2E-06 | 3.8E-05 | NA | NA | NA |
| ENSG00000177943 | *MAMDC4* | 57 | -0.4 | 5.1E-04 | 2.4E-02 | -0.5 | 2.9E-03 | 1.7E-02 | -0.1 | 2.5E-02 | 1.9E-01 |
| ENSG00000140525 | *FANCI* | 581 | 0.3 | 5.1E-04 | 2.4E-02 | 1.0 | 6.4E-16 | 5.7E-14 | 0.1 | 3.6E-02 | 2.3E-01 |
| ENSG00000147852 | *VLDLR* | 2210 | 0.2 | 5.3E-04 | 2.5E-02 | 0.3 | 4.0E-03 | 2.3E-02 | 0.2 | 1.0E-04 | 3.7E-03 |
| ENSG00000127838 | *PNKD* | 2614 | -0.2 | 5.3E-04 | 2.5E-02 | -0.6 | 4.7E-13 | 2.8E-11 | -0.2 | 3.2E-04 | 9.3E-03 |
| ENSG00000129038 | *LOXL1* | 3525 | 0.3 | 5.4E-04 | 2.5E-02 | 1.0 | 1.7E-24 | 3.9E-22 | 0.2 | 2.4E-03 | 4.1E-02 |
| ENSG00000161714 | *PLCD3* | 3289 | -0.2 | 5.5E-04 | 2.5E-02 | -0.3 | 1.8E-05 | 2.4E-04 | -0.2 | 3.1E-03 | 5.0E-02 |
| ENSG00000008735 | *MAPK8IP2* | 416 | -0.3 | 5.5E-04 | 2.5E-02 | -0.6 | 1.6E-03 | 1.1E-02 | -0.2 | 9.4E-03 | 1.1E-01 |
| ENSG00000106348 | *IMPDH1* | 4580 | -0.3 | 5.7E-04 | 2.6E-02 | -0.5 | 6.4E-12 | 3.3E-10 | -0.4 | 1.9E-07 | 1.9E-05 |
| ENSG00000256235 | *SMIM3* | 455 | 0.3 | 5.8E-04 | 2.6E-02 | 0.6 | 1.4E-05 | 1.9E-04 | 0.3 | 1.7E-04 | 5.5E-03 |
| ENSG00000131100 | *ATP6V1E1* | 3005 | 0.1 | 5.8E-04 | 2.6E-02 | 0.2 | 3.2E-02 | 1.1E-01 | 0.1 | 3.5E-03 | 5.4E-02 |
| ENSG00000064666 | *CNN2* | 24511 | -0.1 | 5.8E-04 | 2.7E-02 | -0.2 | 9.3E-03 | 4.5E-02 | -0.1 | 3.9E-03 | 5.8E-02 |
| ENSG00000123352 | *SPATS2* | 537 | 0.2 | 5.9E-04 | 2.7E-02 | 0.3 | 1.9E-03 | 1.2E-02 | 0.1 | 2.0E-02 | 1.6E-01 |
| ENSG00000138760 | *SCARB2* | 4353 | 0.2 | 6.0E-04 | 2.7E-02 | 0.4 | 9.0E-07 | 1.7E-05 | 0.1 | 1.5E-02 | 1.4E-01 |
| ENSG00000134884 | *ARGLU1* | 667 | -0.2 | 6.0E-04 | 2.7E-02 | -0.1 | 1.7E-01 | 3.5E-01 | -0.2 | 2.4E-04 | 7.5E-03 |
| ENSG00000040608 | *RTN4R* | 169 | -0.4 | 6.1E-04 | 2.7E-02 | -1.1 | 6.9E-08 | 1.7E-06 | -0.4 | 1.7E-04 | 5.7E-03 |
| ENSG00000114030 | *KPNA1* | 2590 | -0.1 | 6.1E-04 | 2.7E-02 | -0.2 | 8.2E-03 | 4.0E-02 | -0.1 | 5.4E-03 | 7.2E-02 |
| ENSG00000105447 | *GRWD1* | 1388 | -0.2 | 6.2E-04 | 2.8E-02 | 0.0 | 8.0E-01 | 9.0E-01 | -0.4 | 1.7E-11 | 4.2E-09 |
| ENSG00000197457 | *STMN3* | 1242 | -0.3 | 6.2E-04 | 2.8E-02 | -0.6 | 6.8E-07 | 1.3E-05 | -0.3 | 9.3E-06 | 5.7E-04 |
| ENSG00000111266 | *DUSP16* | 685 | -0.2 | 6.2E-04 | 2.8E-02 | -0.7 | 8.4E-09 | 2.6E-07 | -0.1 | 5.0E-02 | 2.8E-01 |
| ENSG00000124134 | *KCNS1* | 22 | 0.3 | 6.3E-04 | 2.8E-02 | 0.5 | 3.2E-03 | 1.9E-02 | 0.1 | 2.7E-02 | NA |
| ENSG00000132475 | *H3-3B* | 6375 | 0.2 | 6.3E-04 | 2.8E-02 | 0.5 | 3.4E-10 | 1.4E-08 | 0.2 | 4.0E-05 | 1.9E-03 |
| ENSG00000169018 | *FEM1B* | 1942 | -0.2 | 6.4E-04 | 2.8E-02 | -0.3 | 6.3E-04 | 5.0E-03 | -0.1 | 2.9E-03 | 4.7E-02 |
| ENSG00000088727 | *KIF9* | 71 | -0.3 | 6.5E-04 | 2.9E-02 | -0.1 | 2.1E-01 | 4.1E-01 | -0.4 | 3.2E-04 | 9.2E-03 |
| ENSG00000173153 | *ESRRA* | 576 | -0.2 | 6.6E-04 | 2.9E-02 | -0.4 | 2.8E-05 | 3.5E-04 | -0.1 | 3.0E-02 | 2.1E-01 |
| ENSG00000143545 | *RAB13* | 416 | 0.2 | 6.6E-04 | 2.9E-02 | 0.3 | 1.4E-03 | 9.8E-03 | 0.1 | 4.0E-02 | 2.5E-01 |
| ENSG00000154188 | *ANGPT1* | 560 | 0.3 | 6.6E-04 | 2.9E-02 | 1.3 | 1.2E-24 | 2.8E-22 | 0.3 | 1.1E-04 | 3.9E-03 |
| ENSG00000140406 | *TLNRD1* | 1130 | -0.2 | 6.6E-04 | 2.9E-02 | -0.5 | 1.0E-05 | 1.5E-04 | -0.1 | 2.4E-02 | 1.8E-01 |
| ENSG00000094975 | *SUCO* | 634 | 0.2 | 6.7E-04 | 2.9E-02 | 0.2 | 1.8E-02 | 7.2E-02 | 0.3 | 1.3E-04 | 4.4E-03 |
| ENSG00000164323 | *CFAP97* | 745 | 0.3 | 6.8E-04 | 2.9E-02 | 0.3 | 1.1E-02 | 5.1E-02 | 0.4 | 7.1E-05 | 2.8E-03 |
| ENSG00000276023 | *DUSP14* | 2145 | -0.2 | 6.9E-04 | 3.0E-02 | -0.3 | 1.8E-05 | 2.4E-04 | -0.1 | 1.1E-02 | 1.1E-01 |
| ENSG00000106993 | *CDC37L1* | 365 | -0.2 | 6.9E-04 | 3.0E-02 | -0.5 | 1.3E-04 | 1.3E-03 | -0.2 | 8.5E-03 | 9.9E-02 |
| ENSG00000145919 | *BOD1* | 1675 | -0.2 | 7.0E-04 | 3.0E-02 | 0.0 | 7.0E-01 | 8.4E-01 | -0.4 | 8.6E-11 | 1.7E-08 |
| ENSG00000108883 | *EFTUD2* | 3787 | -0.2 | 7.1E-04 | 3.0E-02 | -0.1 | 9.2E-02 | 2.4E-01 | -0.2 | 1.6E-06 | 1.2E-04 |
| ENSG00000177200 | *CHD9* | 556 | 0.2 | 7.1E-04 | 3.0E-02 | 0.3 | 8.0E-04 | 6.1E-03 | 0.2 | 1.6E-03 | 3.1E-02 |
| ENSG00000197217 | *ENTPD4* | 1305 | 0.2 | 7.3E-04 | 3.1E-02 | 0.3 | 1.6E-03 | 1.1E-02 | 0.1 | 4.4E-03 | 6.2E-02 |
| ENSG00000118596 | *SLC16A7* | 346 | 0.2 | 7.4E-04 | 3.1E-02 | 0.6 | 2.0E-07 | 4.4E-06 | 0.1 | 5.0E-02 | 2.8E-01 |
| ENSG00000166503 | *HDGFL3* | 602 | 0.2 | 7.4E-04 | 3.1E-02 | 0.3 | 1.1E-03 | 7.6E-03 | 0.2 | 1.8E-03 | 3.3E-02 |
| ENSG00000174151 | *CYB561D1* | 491 | -0.2 | 7.5E-04 | 3.1E-02 | -0.2 | 4.4E-02 | 1.4E-01 | -0.2 | 1.2E-03 | 2.4E-02 |

| | | | | | | | | | | | |
|---|---|---|---|---|---|---|---|---|---|---|---|
| ENSG00000103888 | *CEMIP* | 15592 | 0.3 | 7.5E-04 | 3.2E-02 | 0.8 | 9.3E-12 | 4.6E-10 | 0.2 | 1.6E-03 | 3.1E-02 |
| ENSG00000103196 | *CRISPLD2* | 3586 | 0.3 | 7.6E-04 | 3.2E-02 | 0.7 | 9.5E-10 | 3.5E-08 | 0.3 | 2.2E-07 | 2.2E-05 |
| ENSG00000286619 | *RP4-681L3.3* | 51 | 0.3 | 7.6E-04 | 3.2E-02 | 0.2 | 1.6E-01 | 3.4E-01 | 0.3 | 1.7E-03 | 3.2E-02 |
| ENSG00000177363 | *LRRN4CL* | 21 | 0.2 | 7.7E-04 | 3.2E-02 | 0.9 | 2.3E-05 | 3.0E-04 | 0.0 | 7.9E-02 | NA |
| ENSG00000049245 | *VAMP3* | 4790 | -0.2 | 7.7E-04 | 3.2E-02 | -0.3 | 6.5E-04 | 5.1E-03 | -0.1 | 1.1E-02 | 1.2E-01 |
| ENSG00000118200 | *CAMSAP2* | 807 | 0.2 | 7.8E-04 | 3.2E-02 | 0.3 | 4.0E-04 | 3.5E-03 | 0.1 | 1.1E-02 | 1.2E-01 |
| ENSG00000111057 | *KRT18* | 40711 | -0.1 | 8.1E-04 | 3.3E-02 | -0.2 | 4.7E-02 | 1.5E-01 | -0.1 | 1.6E-03 | 3.1E-02 |
| ENSG00000141542 | *RAB40B* | 442 | 0.2 | 8.2E-04 | 3.4E-02 | 0.5 | 3.7E-04 | 3.3E-03 | 0.1 | 1.7E-02 | 1.5E-01 |
| ENSG00000164542 | *KIAA0895* | 288 | -0.3 | 8.3E-04 | 3.4E-02 | -1.3 | 1.3E-09 | 4.7E-08 | -0.2 | 6.3E-03 | 8.0E-02 |
| ENSG00000185215 | *TNFAIP2* | 507 | -0.3 | 8.4E-04 | 3.4E-02 | -0.4 | 3.4E-06 | 5.5E-05 | -0.2 | 2.2E-03 | 3.8E-02 |
| ENSG00000100292 | *HMOX1* | 909 | 0.2 | 8.5E-04 | 3.5E-02 | 0.7 | 1.8E-13 | 1.1E-11 | 0.1 | 7.1E-03 | 8.8E-02 |
| ENSG00000180957 | *PITPNB* | 1306 | 0.2 | 8.5E-04 | 3.5E-02 | 0.3 | 9.8E-05 | 1.0E-03 | 0.1 | 5.0E-02 | 2.8E-01 |
| ENSG00000117174 | *ZNHIT6* | 515 | -0.2 | 8.5E-04 | 3.5E-02 | -0.2 | 2.7E-02 | 9.8E-02 | -0.2 | 2.5E-03 | 4.2E-02 |
| ENSG00000164237 | *CMBL* | 1718 | -0.2 | 8.6E-04 | 3.5E-02 | -0.5 | 6.0E-10 | 2.3E-08 | -0.1 | 6.5E-03 | 8.2E-02 |
| ENSG00000087076 | *HSD17B14* | 432 | 0.2 | 8.6E-04 | 3.5E-02 | 0.4 | 1.3E-03 | 9.0E-03 | 0.1 | 9.5E-03 | 1.1E-01 |
| ENSG00000188735 | *TMEM120B* | 465 | 0.2 | 8.7E-04 | 3.5E-02 | 0.1 | 1.3E-01 | 3.0E-01 | 0.2 | 1.1E-03 | 2.4E-02 |
| ENSG00000101935 | *AMMECR1* | 593 | -0.3 | 8.7E-04 | 3.5E-02 | -0.5 | 7.3E-05 | 8.1E-04 | -0.3 | 1.5E-05 | 8.4E-04 |
| ENSG00000206053 | *JPT2* | 3512 | -0.1 | 8.7E-04 | 3.5E-02 | -0.1 | 1.9E-01 | 3.8E-01 | -0.1 | 3.2E-03 | 5.1E-02 |
| ENSG00000088305 | *DNMT3B* | 308 | -0.3 | 8.8E-04 | 3.5E-02 | -0.3 | 2.7E-02 | 9.8E-02 | -0.7 | 5.5E-18 | 2.5E-15 |
| ENSG00000117385 | *P3H1* | 3774 | 0.2 | 8.9E-04 | 3.5E-02 | 0.7 | 3.8E-10 | 1.5E-08 | 0.2 | 4.7E-04 | 1.3E-02 |
| ENSG00000197956 | *S100A6* | 13584 | -0.2 | 8.9E-04 | 3.5E-02 | -0.3 | 7.4E-06 | 1.1E-04 | -0.2 | 7.0E-03 | 8.7E-02 |
| ENSG00000152034 | *MCHR2* | 17 | 0.3 | 9.1E-04 | 3.6E-02 | NA | NA | NA | 0.3 | 1.0E-03 | NA |
| ENSG00000166908 | *PIP4K2C* | 1089 | -0.2 | 9.2E-04 | 3.6E-02 | -0.2 | 4.3E-03 | 2.4E-02 | -0.2 | 1.5E-03 | 3.0E-02 |
| ENSG00000134222 | *PSRC1* | 545 | 0.2 | 9.3E-04 | 3.7E-02 | 0.7 | 1.9E-08 | 5.4E-07 | 0.1 | 6.2E-02 | 3.1E-01 |
| ENSG00000055147 | *FAM114A2* | 514 | 0.2 | 9.5E-04 | 3.7E-02 | 0.0 | 5.8E-01 | 7.6E-01 | 0.3 | 1.4E-04 | 4.7E-03 |
| ENSG00000105866 | *SP4* | 70 | 0.3 | 9.5E-04 | 3.7E-02 | 0.4 | 6.8E-03 | 3.4E-02 | 0.1 | 3.4E-02 | 2.2E-01 |
| ENSG00000171766 | *GATM* | 231 | -0.2 | 9.6E-04 | 3.8E-02 | 0.0 | 9.3E-01 | NA | -0.4 | 3.2E-05 | 1.6E-03 |
| ENSG00000075413 | *MARK3* | 1269 | 0.2 | 9.6E-04 | 3.8E-02 | 0.1 | 1.2E-01 | 2.9E-01 | 0.2 | 1.8E-03 | 3.4E-02 |
| ENSG00000158615 | *PPP1R15B* | 1444 | -0.2 | 9.7E-04 | 3.8E-02 | -0.4 | 1.3E-06 | 2.4E-05 | -0.1 | 1.0E-02 | 1.1E-01 |
| ENSG00000131697 | *NPHP4* | 253 | -0.2 | 9.7E-04 | 3.8E-02 | -0.6 | 4.8E-06 | 7.6E-05 | -0.1 | 3.2E-02 | 2.2E-01 |
| ENSG00000117500 | *TMED5* | 1275 | 0.2 | 9.9E-04 | 3.8E-02 | 0.2 | 1.8E-02 | 7.4E-02 | 0.1 | 1.0E-02 | 1.1E-01 |
| ENSG00000161813 | *LARP4* | 722 | 0.2 | 9.9E-04 | 3.8E-02 | 0.7 | 1.5E-09 | 5.5E-08 | 0.1 | 9.6E-02 | 3.9E-01 |
| ENSG00000066117 | *SMARCD1* | 2595 | -0.2 | 1.0E-03 | 3.9E-02 | -0.4 | 2.3E-08 | 6.4E-07 | -0.2 | 1.1E-04 | 4.0E-03 |
| ENSG00000125864 | *BFSP1* | 128 | 0.2 | 1.0E-03 | 3.9E-02 | 0.3 | 3.4E-02 | 1.2E-01 | 0.1 | 2.7E-02 | 2.0E-01 |
| ENSG00000155016 | *CYP2U1* | 581 | 0.2 | 1.0E-03 | 3.9E-02 | 0.3 | 8.8E-04 | 6.5E-03 | 0.1 | 1.2E-02 | 1.3E-01 |
| ENSG00000152926 | *ZNF117* | 61 | -0.3 | 1.0E-03 | 3.9E-02 | -0.4 | 8.7E-03 | 4.2E-02 | -0.1 | 5.4E-02 | 2.9E-01 |
| ENSG00000255435 | *RP11-770J1.3* | 30 | 0.3 | 1.0E-03 | 3.9E-02 | 0.1 | 4.7E-01 | 6.8E-01 | 0.2 | 3.6E-03 | 5.4E-02 |
| ENSG00000215030 | *RPL13P12* | 749 | -0.2 | 1.1E-03 | 4.0E-02 | -0.2 | 6.0E-02 | 1.8E-01 | -0.3 | 5.7E-04 | 1.4E-02 |
| ENSG00000119231 | *SENP5* | 668 | 0.2 | 1.0E-03 | 4.0E-02 | 0.2 | 1.6E-02 | 6.6E-02 | 0.1 | 1.9E-02 | 1.6E-01 |
| ENSG00000137504 | *CREBZF* | 481 | -0.3 | 1.1E-03 | 4.0E-02 | -0.6 | 1.3E-07 | 3.1E-06 | -0.4 | 2.0E-05 | 1.1E-03 |
| ENSG00000214517 | *PPME1* | 4312 | -0.2 | 1.1E-03 | 4.0E-02 | -0.7 | 1.3E-15 | 1.1E-13 | -0.2 | 3.5E-05 | 1.7E-03 |
| ENSG00000143401 | *ANP32E* | 906 | 0.2 | 1.1E-03 | 4.1E-02 | 0.7 | 4.7E-08 | 1.2E-06 | 0.1 | 3.2E-02 | 2.2E-01 |
| ENSG00000148411 | *NACC2* | 5177 | -0.2 | 1.1E-03 | 4.1E-02 | -0.7 | 5.3E-18 | 5.9E-16 | -0.1 | 5.0E-02 | 2.8E-01 |
| ENSG00000180822 | *PSMG4* | 268 | -0.2 | 1.1E-03 | 4.1E-02 | -0.2 | 4.2E-02 | 1.4E-01 | -0.1 | 2.2E-02 | 1.7E-01 |
| ENSG00000177106 | *EPS8L2* | 1241 | -0.2 | 1.1E-03 | 4.1E-02 | -0.7 | 1.5E-13 | 1.0E-11 | -0.3 | 5.3E-07 | 5.1E-05 |
| ENSG00000106683 | *LIMK1* | 3049 | -0.2 | 1.1E-03 | 4.2E-02 | -0.4 | 1.4E-06 | 2.4E-05 | -0.2 | 3.3E-03 | 5.1E-02 |
| ENSG00000164850 | *GPER1* | 696 | 0.2 | 1.1E-03 | 4.3E-02 | 0.9 | 4.6E-09 | 1.5E-07 | 0.1 | 6.8E-02 | 3.3E-01 |
| ENSG00000146281 | *PM20D2* | 320 | 0.3 | 1.2E-03 | 4.3E-02 | 1.5 | 2.4E-20 | 3.4E-18 | 0.1 | 4.4E-02 | 2.6E-01 |
| ENSG00000276168 | *RN7SL1* | 558 | 0.3 | 1.2E-03 | 4.3E-02 | 0.1 | 6.2E-01 | 7.9E-01 | 0.3 | 3.2E-04 | 9.1E-03 |
| ENSG00000256229 | *ZNF486* | 129 | -0.2 | 1.2E-03 | 4.3E-02 | -0.3 | 2.1E-02 | 8.4E-02 | -0.1 | 2.3E-02 | 1.8E-01 |
| ENSG00000105968 | *H2AZ2* | 3805 | 0.2 | 1.2E-03 | 4.3E-02 | 0.3 | 2.1E-05 | 2.7E-04 | 0.2 | 8.2E-04 | 1.9E-02 |
| ENSG00000143367 | *TUFT1* | 1427 | -0.2 | 1.2E-03 | 4.3E-02 | -0.5 | 1.0E-07 | 2.4E-06 | -0.2 | 5.6E-06 | 3.6E-04 |
| ENSG00000102302 | *FGD1* | 1040 | 0.2 | 1.2E-03 | 4.3E-02 | 0.2 | 2.5E-03 | 1.6E-02 | 0.2 | 6.2E-03 | 7.9E-02 |
| ENSG00000088812 | *ATRN* | 1685 | 0.2 | 1.2E-03 | 4.3E-02 | 0.2 | 1.4E-02 | 5.9E-02 | 0.2 | 6.5E-04 | 1.6E-02 |
| ENSG00000144118 | *RALB* | 2737 | -0.2 | 1.2E-03 | 4.3E-02 | -0.6 | 2.3E-10 | 9.6E-09 | -0.1 | 4.8E-02 | 2.7E-01 |
| ENSG00000100526 | *CDKN3* | 532 | 0.2 | 1.2E-03 | 4.3E-02 | 0.8 | 1.4E-09 | 4.8E-08 | 0.1 | 2.8E-02 | 2.0E-01 |
| ENSG00000135018 | *UBQLN1* | 2619 | 0.2 | 1.2E-03 | 4.4E-02 | 0.6 | 8.4E-13 | 4.9E-11 | 0.1 | 4.0E-02 | 2.5E-01 |
| ENSG00000173218 | *VANGL1* | 991 | 0.2 | 1.2E-03 | 4.4E-02 | 1.0 | 1.9E-19 | 2.4E-17 | 0.1 | 4.4E-03 | 6.3E-02 |
| ENSG00000213585 | *VDAC1* | 5727 | -0.1 | 1.2E-03 | 4.5E-02 | -0.2 | 8.4E-04 | 6.3E-03 | -0.1 | 2.8E-02 | 2.0E-01 |
| ENSG00000103978 | *TMEM87A* | 1086 | 0.3 | 1.2E-03 | 4.5E-02 | 0.9 | 1.0E-17 | 1.1E-15 | 0.5 | 5.2E-11 | 1.1E-08 |
| ENSG00000070778 | *PTPN21* | 1199 | -0.2 | 1.2E-03 | 4.5E-02 | -0.3 | 1.0E-04 | 1.1E-03 | -0.1 | 4.9E-02 | 2.7E-01 |
| ENSG00000175283 | *DOLK* | 945 | 0.2 | 1.3E-03 | 4.5E-02 | 0.7 | 2.8E-08 | 7.5E-07 | 0.1 | 7.7E-03 | 9.3E-02 |
| ENSG00000198205 | *ZXDA* | 79 | 0.3 | 1.3E-03 | 4.5E-02 | 0.3 | 2.8E-02 | 1.0E-01 | 0.1 | 1.8E-02 | 1.5E-01 |
| ENSG00000009780 | *FAM76A* | 229 | 0.2 | 1.3E-03 | 4.5E-02 | 0.1 | 2.8E-01 | 4.9E-01 | 0.2 | 3.0E-03 | 4.8E-02 |
| ENSG00000104522 | *GFUS* | 1916 | -0.1 | 1.3E-03 | 4.5E-02 | -0.1 | 9.6E-02 | 2.4E-01 | -0.1 | 2.0E-02 | 1.6E-01 |
| ENSG00000152242 | *C18orf25* | 725 | -0.1 | 1.3E-03 | 4.6E-02 | -0.2 | 4.5E-02 | 1.4E-01 | -0.1 | 1.3E-02 | 1.3E-01 |
| ENSG00000171462 | *DLK2* | 123 | -0.3 | 1.3E-03 | 4.6E-02 | -0.8 | 9.7E-08 | 2.3E-06 | -0.2 | 2.9E-03 | 4.7E-02 |
| ENSG00000166128 | *RAB8B* | 819 | -0.2 | 1.3E-03 | 4.6E-02 | -0.2 | 2.3E-02 | 8.9E-02 | -0.2 | 9.6E-04 | 2.1E-02 |
| ENSG00000137710 | *RDX* | 2136 | 0.2 | 1.3E-03 | 4.7E-02 | 0.2 | 3.7E-02 | 1.2E-01 | 0.2 | 1.8E-03 | 3.3E-02 |
| ENSG00000106105 | *GARS1* | 7956 | -0.2 | 1.3E-03 | 4.7E-02 | -0.3 | 2.3E-05 | 2.9E-04 | -0.1 | 2.1E-03 | 3.7E-02 |
| ENSG00000133028 | *SCO1* | 1185 | -0.2 | 1.3E-03 | 4.7E-02 | -0.3 | 2.0E-04 | 1.9E-03 | -0.1 | 3.1E-02 | 2.1E-01 |
| ENSG00000158008 | *EXTL1* | 73 | 0.3 | 1.4E-03 | 4.8E-02 | 0.8 | 1.0E-06 | 1.9E-05 | 0.0 | 3.1E-01 | 6.6E-01 |
| ENSG00000166938 | *DIS3L* | 607 | 0.2 | 1.4E-03 | 4.9E-02 | 0.3 | 4.4E-03 | 2.5E-02 | 0.1 | 2.5E-02 | 1.9E-01 |
| ENSG00000124789 | *NUP153* | 1535 | 0.2 | 1.4E-03 | 4.9E-02 | 0.3 | 5.2E-04 | 4.3E-03 | 0.1 | 1.8E-02 | 1.5E-01 |

**Supplementary Table 6**

| Ontology | ID | Description | Gene Ratio | Background Ratio | Enrichment | P | P.adj | Gene names |
|---|---|---|---|---|---|---|---|---|
| BP | GO:0016236 | macroautophagy | 23/428 | 350/23153 | 3.6 | 1.8E-07 | 0.0009 | *VDAC1/UBQLN1/RALB/PIP4K2C/HMOX1/ATP6V1E1/ATG101/VPS4B/TP53/SCOC/ATG16L1/ATP6V0A2/STX12/TMEM41B/ERN1/TP53INP1/MAP3K7/LIX1L/PIP4K2B/RAB7A/ATG5/UBQLN2/NBR1* |
| BP | GO:0000045 | autophagosome assembly | 12/428 | 128/23153 | 5.1 | 4.7E-06 | 0.0078 | *UBQLN1/RALB/PIP4K2C/ATG101/ATG16L1/STX12/TMEM41B/TP53INP1/PIP4K2B/RAB7A/ATG5/UBQLN2* |
| CC | GO:0000407 | phagophore assembly site | 7/428 | 39/23153 | 9.7 | 6.5E-06 | 0.0078 | *ATG101/ATG16L1/STX12/BCAS3/RAB7A/ATG5/NBR1* |
| BP | GO:0016241 | regulation of macroautophagy | 14/428 | 182/23153 | 4.2 | 7.9E-06 | 0.0078 | *VDAC1/UBQLN1/RALB/PIP4K2C/HMOX1/ATP6V1E1/TP53/SCOC/ATP6V0A2/ERN1/MAP3K7/PIP4K2B/ATG5/UBQLN2* |
| BP | GO:0007033 | vacuole organization | 16/428 | 235/23153 | 3.7 | 8.6E-06 | 0.0078 | *UBQLN1/RALB/PIP4K2C/SCARB2/ATG101/ATG16L1/CLN6/STX12/CCDC115/TMEM41B/LAMTOR1/TP53INP1/PIP4K2B/RAB7A/ATG5/UBQLN2* |
| BP | GO:1905037 | autophagosome organization | 12/428 | 137/23153 | 4.7 | 9.5E-06 | 0.0078 | *UBQLN1/RALB/PIP4K2C/ATG101/ATG16L1/STX12/TMEM41B/TP53INP1/PIP4K2B/RAB7A/ATG5/UBQLN2* |
| BP | GO:0032869 | cellular response to insulin stimulus | 15/428 | 218/23153 | 3.7 | 1.4E-05 | 0.0092 | *PIP4K2C/RAB13/ESRRA/SLC39A14/ZDHHC7/ZNF106/NDEL1/ENPP1/PIK3R3/YWHAG/ERRFI1/PIP4K2B/GRB10/LPIN2/PCK2* |
| BP | GO:0007265 | Ras protein signal transduction | 21/428 | 392/23153 | 2.9 | 1.5E-05 | 0.0092 | *RDX/RALB/LIMK1/EPS8L2/STMN3/RTN4R/CYTH1/TP53/GIT2/F2R/KCTD10/CDC42SE1/HACD3/NKIRAS2/RIT1/FARP2/MAPKAP1/RASGEF1A/CDC42SE2/RAP2B/RAPGEF2* |
